# Nm-Nano: A Machine Learning Framework for Transcriptome-Wide Single Molecule Mapping of 2´-O-Methylation (Nm) Sites in Nanopore Direct RNA Sequencing Datasets

**DOI:** 10.1101/2022.01.03.473214

**Authors:** Doaa Hassan, Aditya Ariyur, Swapna Vidhur Daulatabad, Quoseena Mir, Sarath Chandra Janga

**Author notes:** Correspondence (S.C.J.).

## Abstract

Nm (2’-O-methylation) is one of the most abundant modifications of mRNAs and non-coding RNAs. It has a great contribution in many biological processes such as the normal functioning of tRNA, the protection of mRNA against degradation by DXO protein, and the biogenesis and specificity of rRNA. Recently, the single-molecule sequencing techniques for long reads of RNA sequences data offered by Oxford Nanopore technologies have enabled the direct detection of RNA modifications on the molecule that is being sequenced. In this paper, we propose a bio-computational framework, Nm-Nano for predicting the existence of Nm sites in Nanopore direct RNA sequencing reads of human cell lines. This addresses the limitations of Nm predictors presented in the literature that were only able to detect those sites on short reads of RNA sequences data of cell lines of different species or long read sequencing data of non-human cell lines (yeast). Nm-Nano framework integrates two supervised machine learning (ML) models for predicting Nm sites in Nanopore direct RNA sequencing data, namely the Extreme Gradient Boosting (XGBoost) and Random Forest (RF) with K-mers embedding models. XGBoost is trained with the features extracted from the modified and unmodified Nanopore signals and their corresponding K-mers resulting from the reported underlying RNA sequence obtained by base-calling, while RF model is trained with the same set of features used to train XGBoost, in addition to a dense vector representation of RNA K-mers generated by word2vec technique. Results on benchmark data sets from Hela and Hek293 cell lines demonstrate high accuracy (99% with XGBoost and 92% with RF) in identifying Nm sites. Deploying Nm-Nano on Hela and Hek293 cell lines reveals the frequently Nm-modified genes. In Hela cell lines, 125 genes are identified as frequently Nm-modified, showing enrichment in ontologies related to immune response and cellular processes. In Hek293 cell lines, 61 genes are identified as frequently Nm-modified, with enrichment in processes like glycolysis and protein localization. These findings underscore the diverse regulatory roles of Nm modifications in metabolic pathways, protein degradation, and cellular processes. The source code of Nm-Nano can be freely accessed at https://github.com/Janga-Lab/Nm-Nano.

## 1. Introduction

The 2’-O-methylation (or Nm, where N denotes any nucleotide) is a co- or post-transcriptional modification of RNA, occurring when a methyl group (–CH3) is added to the 2’ hydroxyl (–OH) of the ribose moiety. This modification can appear on any nucleotide regardless of the type of nitrogenous base. Nm is an abundant modification that occurs frequently in mRNAs and at multiple locations in non-coding RNAs such as transfer RNA (tRNA), ribosomal RNA (rRNA), small nuclear RNA (snRNA) and piRNA [1–4]. This is due to the role that internal 2′-O-methylation of mRNA plays as a new mechanism of genetic regulatory control, with the ability to influence mRNA abundance and protein levels both in vitro and in vivo [5].

Nm modification has a great contribution in many biological processes such as the normal functioning of tRNA [6], protecting mRNA from degradation by DXO protein [7], and the biogenesis and specificity of rRNA [8, 9]. It has been also found that Nm modification has been associated with many human diseases (e.g., cancer and autoimmune diseases) and has potential indirect links to some other biological defects [10].

Detecting Nm modifications in RNAs has been a great challenge for many years and various experimental methods for identifying such modification have been presented in the literature [10]. However, each of these methods has exhibited significant limitations. For example, RiboMethseq [11, 12] was introduced as a sequencing-based method for mapping and quantitation of Nm modification based on simple chemical principle, namely the several orders of magnitude difference in nucleophilicity of a 2′-OH and a 2′-O-Me. It uses the proprietary ligation protocol for direct ligation to 5′-OH and 3′-P ends with alkaline fragmentation to prepare RNA for sequencing. The read-ends of library fragments are used for mapping with nucleotide resolution and calculation of the fraction of molecules methylated at the Nm sites. However, the relative inefficiency of the ligation protocol imposes substantial amounts of input RNA (>1 µg) which requires increasing the sequencing depth. Thus, to address this limitation, another chemical method called RibOxi-seq was presented for detecting Nm modifications in RNAs [13]. Using this method, Nm sites could be mapped after ligation of linkers to the Nm-modified nucleotide at the 3′-end. However this method was only able to identify significantly fewer Nm modification sites relative to those reported by LC-MS/MS methods, a biochemical method to detect and quantify the relative abundance of RNA modification [14, 15]. Despite LC-MS/MS providing industry standard results, it is time and labor consuming, as well as requiring large amounts of input RNA and is limited for low-abundance nucleotides [16]. Recently, Dai et al. introduced a sensitive high-throughput experimental method called Nm-seq which was able to detect Nm sites at low stoichiometry especially in mRNAs with single-base resolution, achieving an outstanding detection of Nm modification [17].

However, in general the experimental methods are naturally costly due to the high labor effort. Therefore, there have been relatively few computational biology methods proposed in the literature to overcome the limitations of experimental methods for detecting RNA Nm modifications [18–20]. Those computational methods mainly relied on developing machine/deep learning classification algorithms to identify Nm sites in RNA sequences based only on short read data and were not applied to long reads with a capacity to sequence on average over 10 kb in one single read, and thereby requiring less reads to cover the same gene. For instance, a support vector machine (SVM)-based method was presented in [18] to identify Nm sites in RNA short reads sequences of the human genome by encoding RNA sequences using nucleotide chemical properties and nucleotide compositions. This model was validated by identifying Nm sites in Mus musculus and Saccharomyces cerevisiae genomes. Another research work presented in [19] proposed a deep learning-based method for identifying Nm sites in short reads RNA sequences. In this approach, dna2vec-a biological sequence embedding method originally inspired by the word2vec model of text analysis was adopted to yield embedded representations of RNA sequences that may or may not contain Nm sites. Those embedded representations were fed as features to a Convolutional Neural Network (CNN) for classification of RNA sequences into modified with Nm sites or not modified. The method was trained using the data collected from Nm-seq experimental method. Another prediction model based on Random Forest for identifying Nm sites in short read RNA sequences was presented in [20]. This model was trained with features extracted by multi-encoding scheme combination that combines the one-hot encoding, together with position-specific dinucleotide sequence profile and K-nucleotide frequency encoding.

Recently, the third-generation sequencing technologies such as the platforms provided by Oxford Nanopore Technologies (ONT) has been proposed as a new means to detect RNA modifications on long RNA sequence data [21]. However, to our knowledge, this technology has been only applied in two studies [22, 23] for detecting Nm modifications. In [22], the main goal was to predict the stoichiometry of Nm-modified sites in yeast mitochondrial rRNA using 2-class (Nm-modified or unmodified) classification algorithms deployed in a tool called nanoRMS [22]. This tool used the characteristic base-calling “error” signatures in the Nanopore data as features for training a supervised or unsupervised learning models to identify the stoichiometry of Nm sites using a threshold for base mismatch frequency in different types of RNAs in yeast. However, nanoRMS was not applied to predict Nm sites in the RNA sequence of human cell lines which are larger and more complex than yeast. Also, the single read features used to train the predictors of nanoRMS were averaged before Nm prediction making it not feasible to obtain the contribution of each feature in predicting Nm sites. Moreover, relying on base-calling errors for detecting RNA modification as in nanoRMS implementation might decrease with the advances of developing high accurate Nanopore base-calling algorithms. In [23], a dual-path framework called HybridNm was proposed to predict Nm subtypes in one human cell line (Hek293) based on features extracted from RNA short reads sequenced with Illumina and RNA long read sequenced with ONT to improve the prediction of Nm sites. Therefore, this framework was not purely relying on ONT technology for predicting Nm sites in RNA sequences. Moreover, the base-calling errors were used as features to distinguish Nm from unmodified sites which again might cause decreasing in the performance of accurately predicting Nm sites with the advances of developing high accurate Nanopore base-calling algorithms. To this end, our work aims to extend this research direction and address nanoRMS and HybridNm limitations by combining ML and ONT Technology to identify Nm sites in long RNA sequence reads of human cell lines based on features extracted from raw Nanopore signals. We have developed a framework called Nm-Nano that integrates two different supervised ML models (predictors) to identify Nm sites in Nanopore direct RNA sequencing reads of Hela and Hek293 cell lines, namely the XGBoost and RF with K-mer embedding models (Figure 1.A and B). The developed predictors integrated in Nm-Nano framework for identifying Nm sites have been trained and tested upon a set of Nm-modified and unmodified Nanopore signals. Those signals were generated by passing a set of ‘modified’ RNA sequences containing Nm sites at known positions (identified using the standard Nm-seq experimental method [17]) and ‘unmodified’ ones through the ONT MinION device.

**Figure1.**
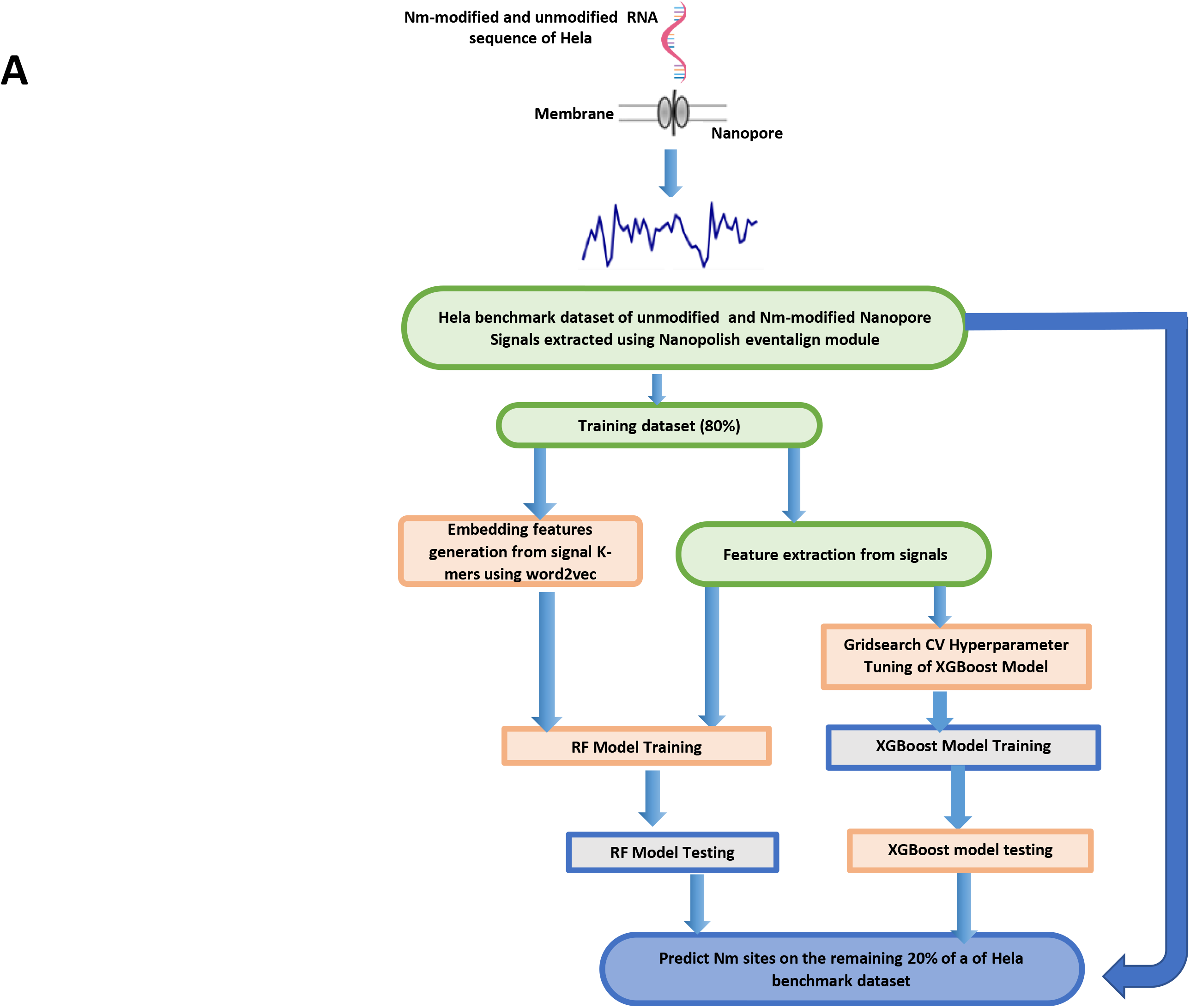
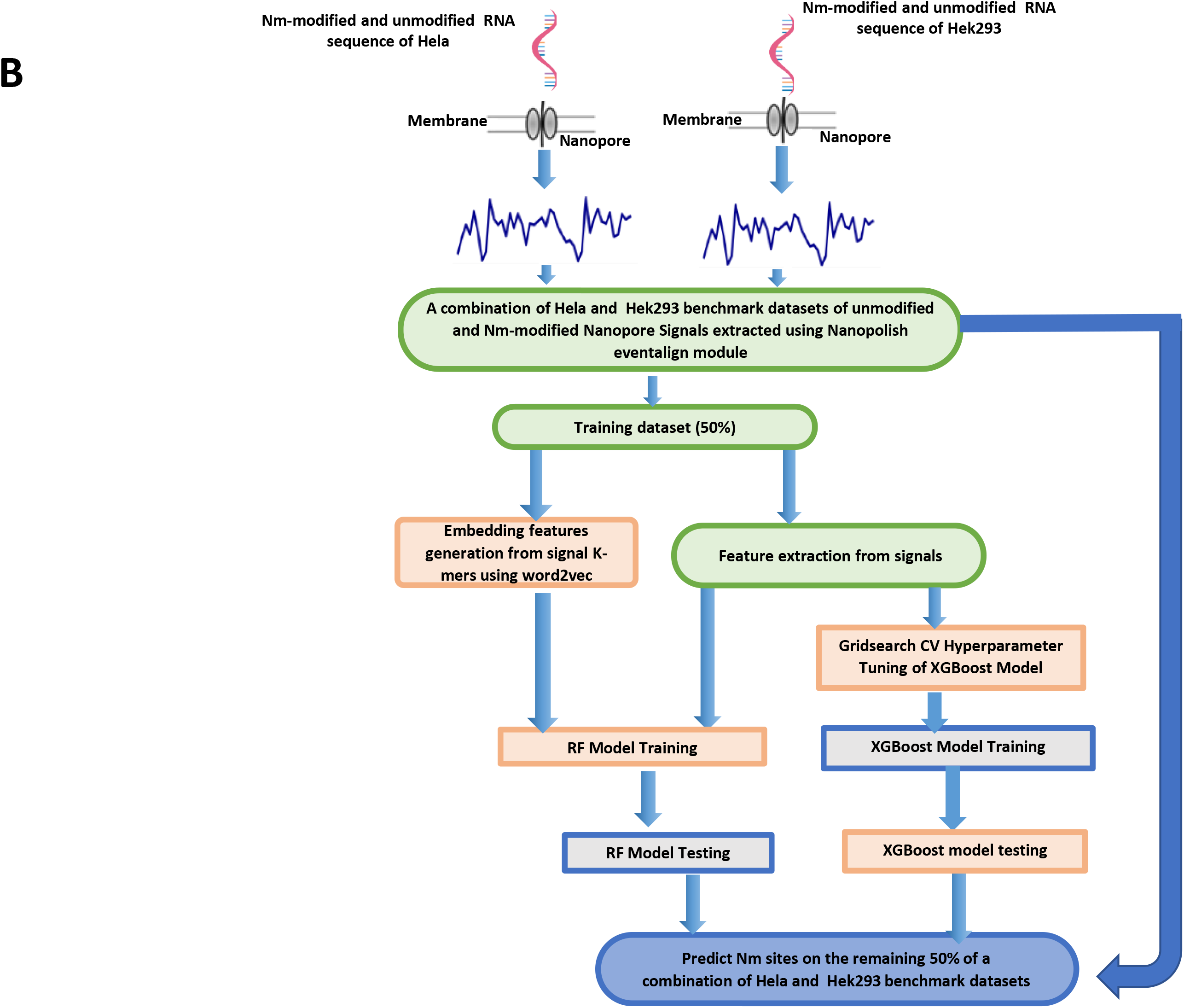
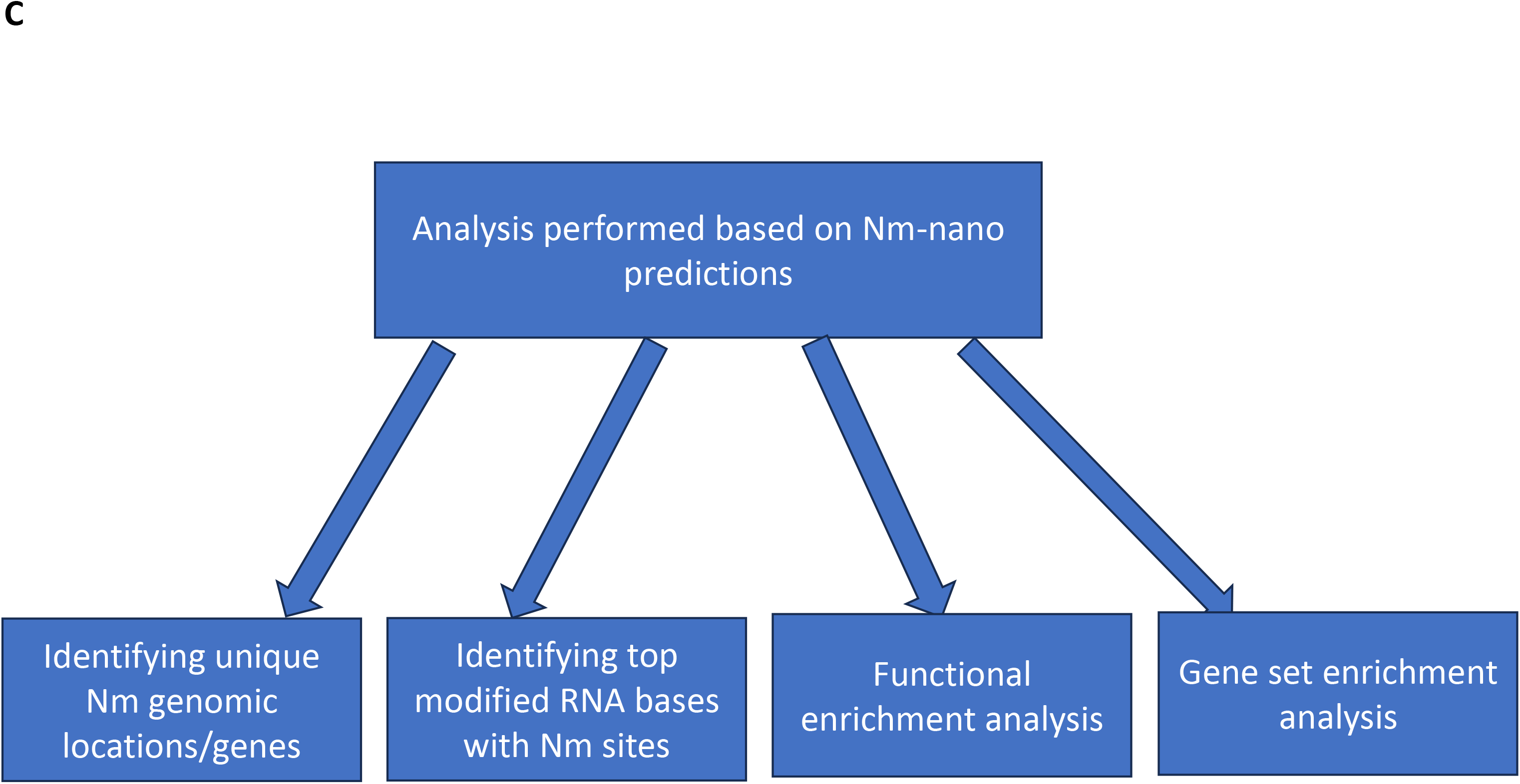
The Nm-Nano framework for predicting Nm sites on (a) Hela cell line using random 80/20 train/test split (b) 50% of the combination of Hela and Hek293 benchmark dataset using integrated validation testing with random 50/50 train/test split on this combination (c) Analysis performed based on Nm-Nano predictions.

By deploying Nm-Nano to predict Nm sites in Nanopore direct RNA sequencing reads of Hela and Hek293 cell lines, we were able to perform various types of biological analysis (Figure 1.C) including: identifying unique Nm genomic locations/genes, identifying top modified RNA bases with Nm sites, functional and gene set enrichment analysis of identified Nm-genes in both cell lines.

## 2. Results

We have used two validation methods when evaluating the performance of Nm-Nano predictors: namely the random-test splitting and integrated validation testing. In the former, the benchmark dataset of Hela cell line introduced in subsection 4.2 in Methods section is randomly divided into two folds: one for training and another for testing. The test size parameter for this method was set to 0.2 which means 80% of the benchmark dataset is used for training the Nm-Nano ML model and 20% of the dataset is kept for testing it. In the latter, a combination of two benchmark datasets for two different cell lines (Hela and Hel293) was used, where 50% of this combination is used for training the Nm-Nano ML model and 50% is kept for testing it.

### 2.1 Performance evaluation with random-test splitting

Table 1 shows the performance of XGBoost and RF with K-mer embedding ML models implemented in Nm-Nano framework that are available on its GitHub page when applied to the Hela benchmark dataset. As the table shows, both models perform very well in detecting Nm sites. However, the XGBoost model outperforms the RF with K-mer embedding in terms of accuracy, precision, recall and Area Under the Curve (AUC).

**Table 1.**
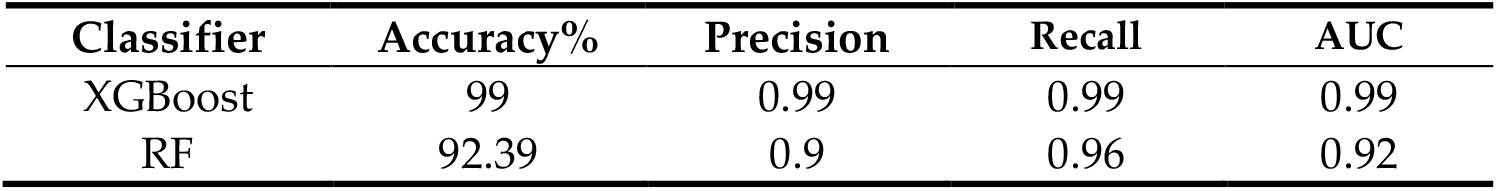
The performance of Nm-nano predictors on Hela benchmark dataset with random-test splitting.

The learning (Figure 2. panels A, and D), and loss (Figure 2. panels B, and E) curves of XGBoost, and RF with K-mer embedding respectively show that the performance of XGBoost in terms of accuracy score and misclassification error outperforms the performance of RF with K-mer embedding. Also, the receiver operating characteristic (ROC) curves of XGBoost and RF with K-mer embedding (Figure 2. panels C, and F respectively) show that the percentage of true positive rate to the false positive rate in case of XGBoost model is more than the one for RF with K-mer embedding model. Supplementary file 1_test_split_Hek293_results.docx and Figure 1_test_split_hek show the performance of XGBoost and RF with K-mer embedding ML models, with random test split on Hek293 benchmark dataset.

**Figure 2.**
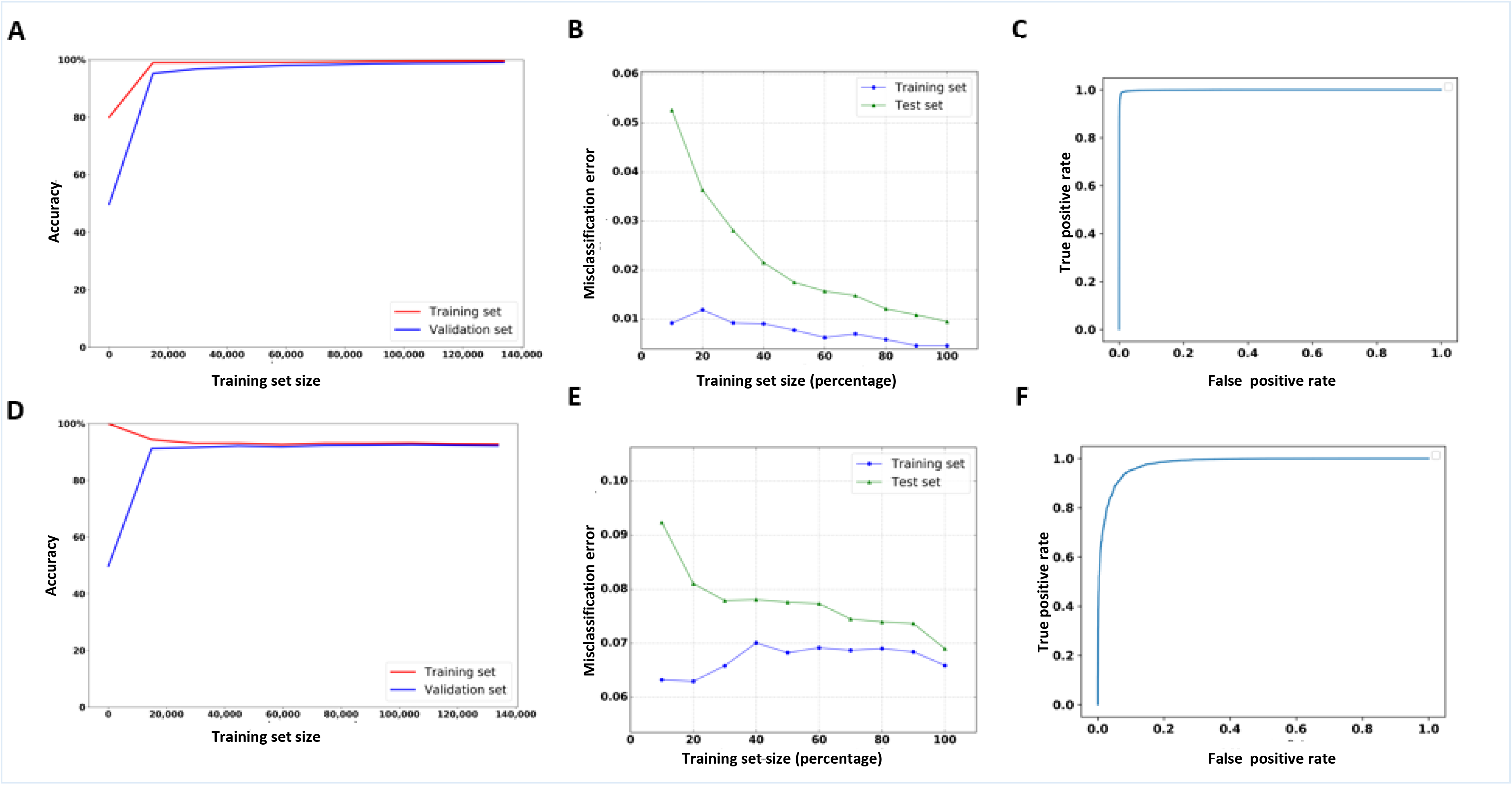
The learning, loss and ROC curves of Nm-nano predictors validated on Hela benchmark dataset with random split testing, where 80% of data is used for training and the remaining 20% is kept for testing. (a, b, and c) XGBoost model and (d, e, and f) RF with K-mer embedding model.

Table 2 shows the performance of Nm-Nano ML models with random test-splitting on Hela benchmark dataset in terms of accuracy with each of the extracted features as well as the embedding features generated using word2vec embedding technique introduced early in Section 1 and later in Subsection 4.4. Clearly the position feature contributes more to the classifiers’ accuracy than other extracted features used for training either the XGBoost or RF with embedding ML models. It is followed by the model mean, then K-mer match features in case of XGboost and K-mer match then model mean feature in case of RF with K-mer embedding. It was also observed that the event/signal standard deviation (Event_stdv) feature achieves the lowest contribution to the performances of XGBoost and RF with embedding models. Table 2 also shows that the embedding features generated by word2vec technique strongly contribute to the performance of RF as those features follow the most contributing feature (i.e., position). Despite the success of these features in improving the performance of RF, they were not used to train the XGBoost model. This is because this model achieved high performance by tuning its parameters with grid search algorithm [24] which takes considerable time to obtain the best values for the parameters.

**Table 2.**
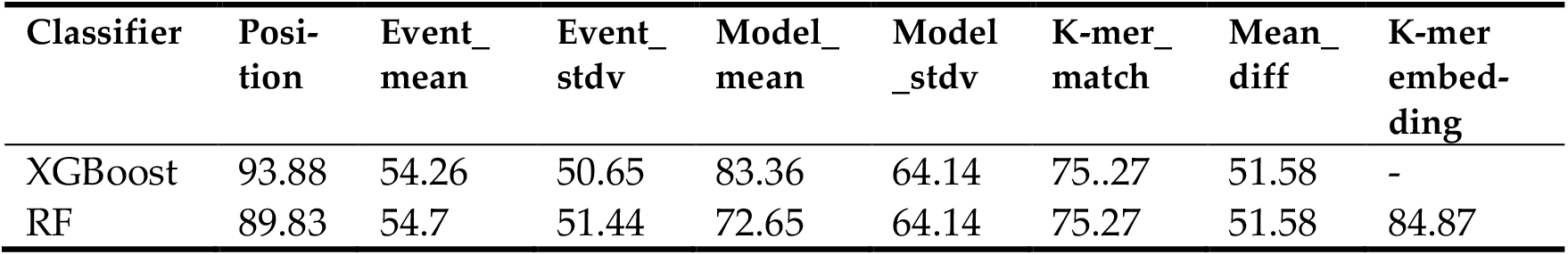
The performance of Nm-Nano predictors on Hela benchmark dataset in terms of accuracy (%) with random test-splitting using single type of feature.

| Classifier | Position | Event_mean | Event_stdv | Model_mean | Model_stdv | K-mer_match | Mean_diff | K-mer_embedding |
| --- | --- | --- | --- | --- | --- | --- | --- | --- |
| XGBoost | 93.88 | 54.26 | 50.65 | 83.36 | 64.14 | 75.27 | 51.58 | - |
| RF | 89.83 | 54.7 | 51.44 | 72.65 | 64.14 | 75.27 | 51.58 | 84.87 |

Therefore, generating more features with word2vec embedding techniques for training the grid-search XGBoost model will add extra processing overhead. This is due to adding the time taken by word2vec technique for generating embedding features to the time taken by the grid search algorithm for hyper parameter tuning of XGBoost model. This will make XGBoost very slow when applying it to the benchmark dataset of a given cell line with a slight improvement in its performance that would not be proportional to the huge increase in the processing time of XGBoost. Table 3 shows the performance of XGBoost (when the model is trained with the extracted features only) versus the performance of XGBoost with K-mer embedding (i.e., when the model is trained with the combination of the extracted features and the embedding reference K-mers features) as well as the execution time in seconds in each case.

**Table 3.** The performance of XGBoost versus the performance of the XGBoost with K-mer embedding model applied to Hela benchmark dataset with random test-splitting.

| Classifier | Accuracy | Precision | Recall | AUC | Execution time (Secs) |
| --- | --- | --- | --- | --- | --- |
| XGBoost | 99 | 0.99 | 0.99 | 0.99 | 43.81 |
| XGBoost with K-mer embedding | 99 | 0.99 | 1 | 0.994 | 608.2 |

### 2.2 Performance evaluation with integrated validation testing

Table 4 shows the performance of ML models with integrated validation testing, where Nm-Nano’s predictors are applied on 50% of combination of Hela and Hek293 benchmark datasets in training phase and tested on the remaining 50% of this combination in the testing phase. As the results show, both models perform very well in predicting Nm sites, though XGBoost model outperforms RF with K-mer embedding model. The learning (Figure 3. panels A, and D), and loss (Figure 3. panels B, and E) curves of XGBoost, and RF with K-mer embedding respectively show that the performance of XGBoost in terms of accuracy score and misclassification error outperforms the performance of RF with K-mer embedding. The receiver operating characteristic (ROC) curves of XGBoost and RF with embedding (Figure 3. panels C, and E respectively) show that the percentage of true positive rate to the false positive rate in case of XGBoost model is more than the one for RF with embedding model.

**Figure 3.**
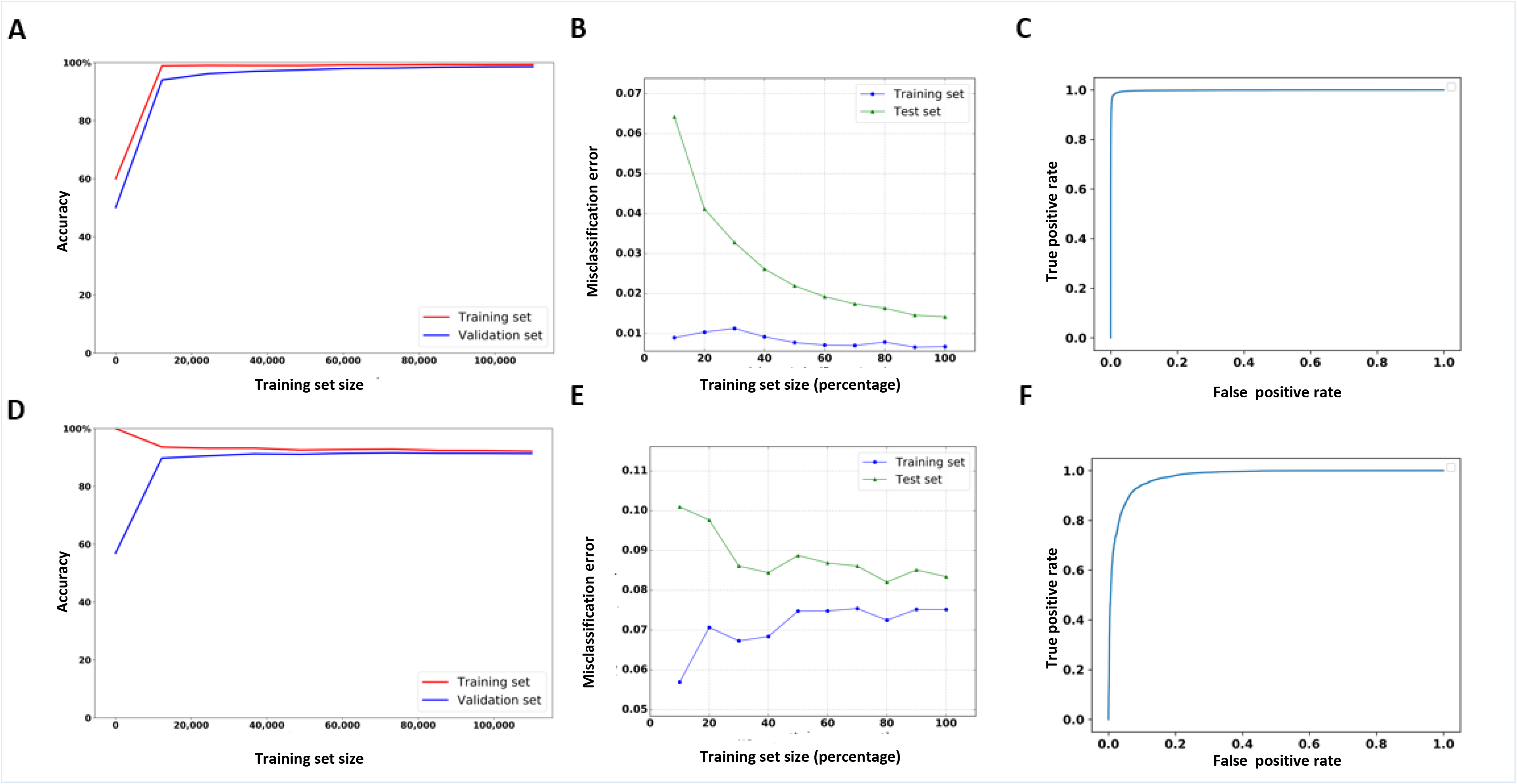
The learning, loss and ROC curves of Nm-nano predictors in integrated validation testing, where 50% of combination of Hela and Hek293 benchmark datasets is used for training and the remaining 50% is used for testing. (a,b, and c) XGBoost model and (d, e, and f) RF with K-mer embedding model.

**Table 4.** The performance of Nm-nano predictors on a combination of Hela & Hek293 benchmark datasets with 0.5 random-test splitting.

| Classifier | Accuracy% | Precision | Recall | AUC |
| --- | --- | --- | --- | --- |
| XGBoost | 98.58 | 0.99 | 0.99 | 0.99 |
| RF | 91.63 | 0.89 | 0.96 | 0.92 |

Table 5 shows the performance of ML models with integrated validation testing in terms of accuracy with single type of feature. This was achieved by testing the performance of Nm-nano predictors with each of the extracted features as well as the embedding features generated using word2vec embedding technique. Clearly the features generated with word2vec embedding technique strongly contribute to the RF classifier accuracy as those features follow the most contributing feature (i.e., position), but they were not considered for training the grid search XGBoost model. Again, this is due to the extra processing overhead resulting from combining the time taken for generating embedding features by wor2vec technique and the time taken by grid search algorithm for obtaining the best parameters values of XGBoost as we early mentioned in subsection 2.1. As for the contribution of each of the seven extracted features, it was observed that the position feature achieves the best among all extracted features followed by model mean feature, then the K-mer match feature for XGBoosat and K-mer match then model mean feature for RF with K-mer embedding. Also, it was observed that event/signal standard deviation (event _stdv) has the lowest contribution to the performance of either XGBoost or RF with K-mer embedding models.

**Table 5.** The performance of Nm-nano predictors with integrated validation testing in terms of accuracy (%) using single type of feature.

| Classifier | Position | Event_mean | Event_stdv | Model_mean | Model_stdv | K-mer_match | Mean_diff | K-mer_embedding |
| --- | --- | --- | --- | --- | --- | --- | --- | --- |
| XGBoost | 94.62 | 53.41 | 51.31 | 81.22 | 62.24 | 75.14 | 51.69 | - |
| RF | 85.45 | 54.19 | 51.71 | 72.54 | 62.24 | 75.14 | 51.7 | 82.92 |

### 2.3. Abundance of Nm sites

In order to identify the abundance of Nm sites in the RNA sequence of either Hela or Hek293 cell lines, first we run XGBoost model (since it outperforms RF with K-mer embedding model) on the complete RNA sequence reads of Hela and Hek293 cell lines. Next, we identify all samples with predicted Nm sites in those reads, then we identify the number of Nm unique genomic locations corresponding to those Nm predictions as well as their frequencies in both cell lines. We found that there are 11,651,518 Nanopore signal samples predicted as samples with Nm sites from a total of 920,643,073 Nanopore signal samples that represent the complete Hela cell line with 1,674,369 unique genomic locations of Nm (Supplementary Table 1_Nm_unique_genomic_locations _hela.xlsx). Similarly, we found that there are 1712344 Nanopore signal samples predicted as samples with Nm sites from a total of 275,056,668 samples that present the complete RNA sequence of Hek293 cell line with 291,382 unique genomic locations of Nm modification (Supplementary Table 2_Nm_unique_genomic_locations _hek.xlsx). The reference K-mers corresponding to modified Nanopore signals with Nm predictions in Hela and Hek293 cell lines can be identified as strong K-mers in comparison with the reference K-mers corresponding to the unmodified/control Nanopore signals, which can be considered as weak contributors to the Nm prediction. The frequency of those strong reference K-mers that provide an overview of their abundance and show their contribution to Nm predictions in Hela and Hek293 cells lines are available in Supplementary Tables 3_Nm_unique_reference_kmer_freq_hela.xlsx and 4_Nm_unique_reference_kmer_freq_hek.xls respectively. Also, Supplementary Figures 2_top_10-modified_bases_hela and 3_top_10-modified_bases_hek provide the sequence logo for the top ten modified bases corresponding to Nm prediction in Hela and Hek293 cell lines respectively.

We found that there are 105678 modified genomic locations common between Hela and Hek293 cell lines (Figure 4.A). Also, we observed that there were 10 genes shared across the top 1% of Nm Modified genes in Hela and Hek293 cell lines (Figures 4.B). Clearly, we notice that the extent of Nm modification (the number of Nanopore signal samples predicted as samples with Nm sites to the total number of Nanopore signal samples either modified with Nm or not modified) in RNA sequences of Hela cell line is higher than its counterpart in Hek293 cell line (1.27 % for Hela versus 0.62% for Hek293). Therefore, the distribution of Nm across normalized gene length for Hela cell line is higher than its equivalent in Hek293 cell line (Figures 4.C). Additionally, and as a primary observation of Figure 4C, we found that Nm modifications are likely to be more prevalent in the 3’ region compared to the 5’ when observed at a transcriptomic level. This distribution reinforces our previous observation about (psuedouridine) RNA modifications having a preference for 3’ over 5’ [25].

**Figure 4.**
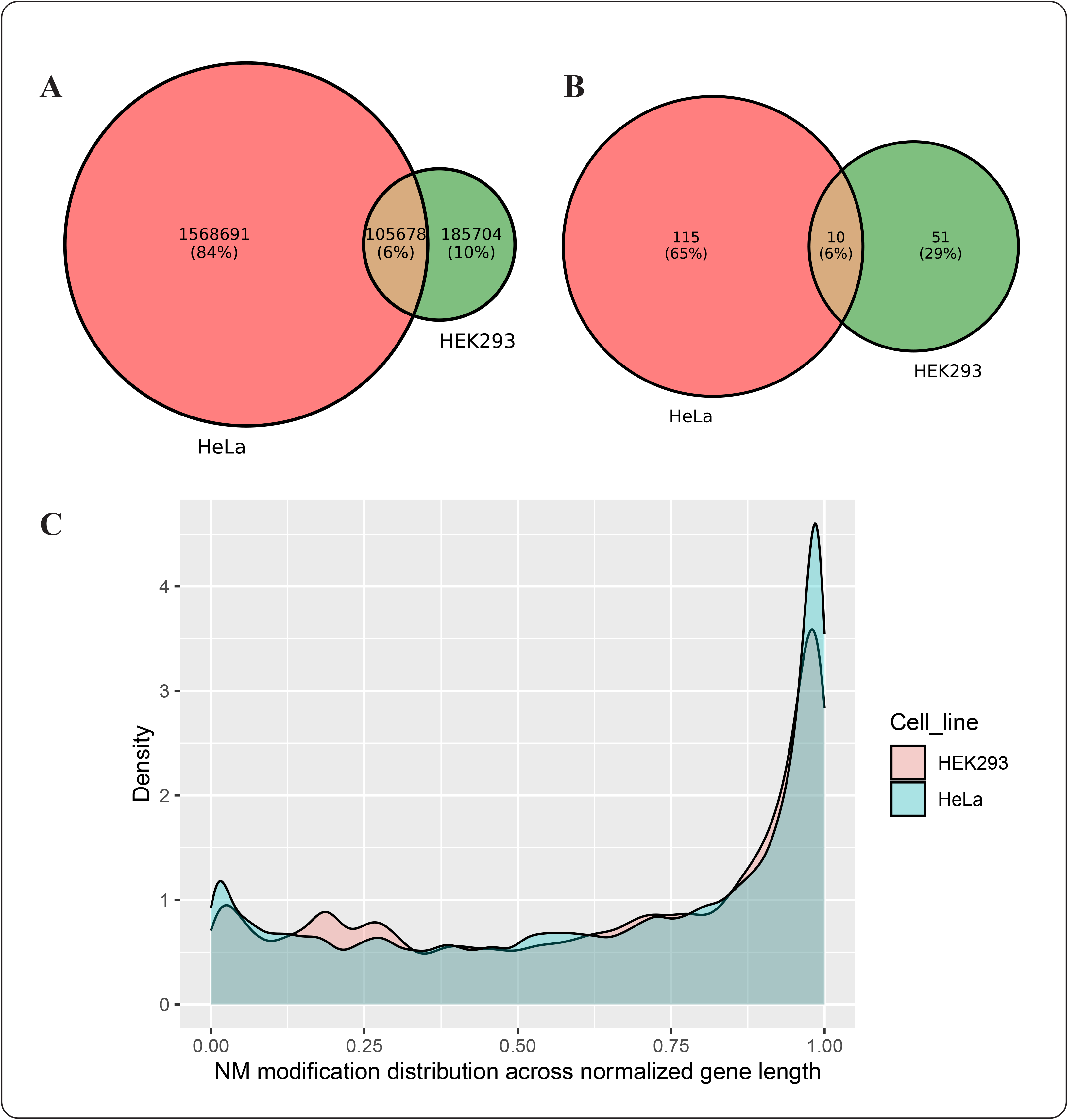
(a) The overlap between Nm unique locations in complete Hek293 and Hela cell lines (b) the overlapping between top frequent 1 % modified Nm genes in complete Hek293 and Hela cell lines (c) The density plots that represents Nm modifications across normalized gene length for Hek293 and Hela cell lines.

Since Nm modifications can occur at any RNA base, we have also reported about the percentage of unique Nm locations occurring per each of the four RNA bases in the two complete cell lines of Hela and Hek293 (Table 6.)

**Table 6.** The percentage of unique Nm locations occurring per each of the four RNA bases in Hela and Hek293 cell lines.

| Cell line | A base | C base | G base | U base |
| --- | --- | --- | --- | --- |
| Hela | 29.75 | 21.54 | 22.91 | 25.81 |
| Hek293 | 26.18% | 25.14% | 26.12% | 22.56% |

### 2.4 Functional enrichment analysis

A total of 61 genes from Hek293 and 125 genes from Hela cell lines were identified as the top 1% frequently modified Nm genes with the highest abundance of Nm modification. The short-listed genes from both cell lines were then plugged into Cytsoscape ClueGo [26] application to obtain the enriched ontologies and pathways at high confidence (p<0.05). Enrichment observations from this analysis are visualized in Figure 5 A, and B for Hek293 and Hela cell lines respectively.

**Figure 5.**
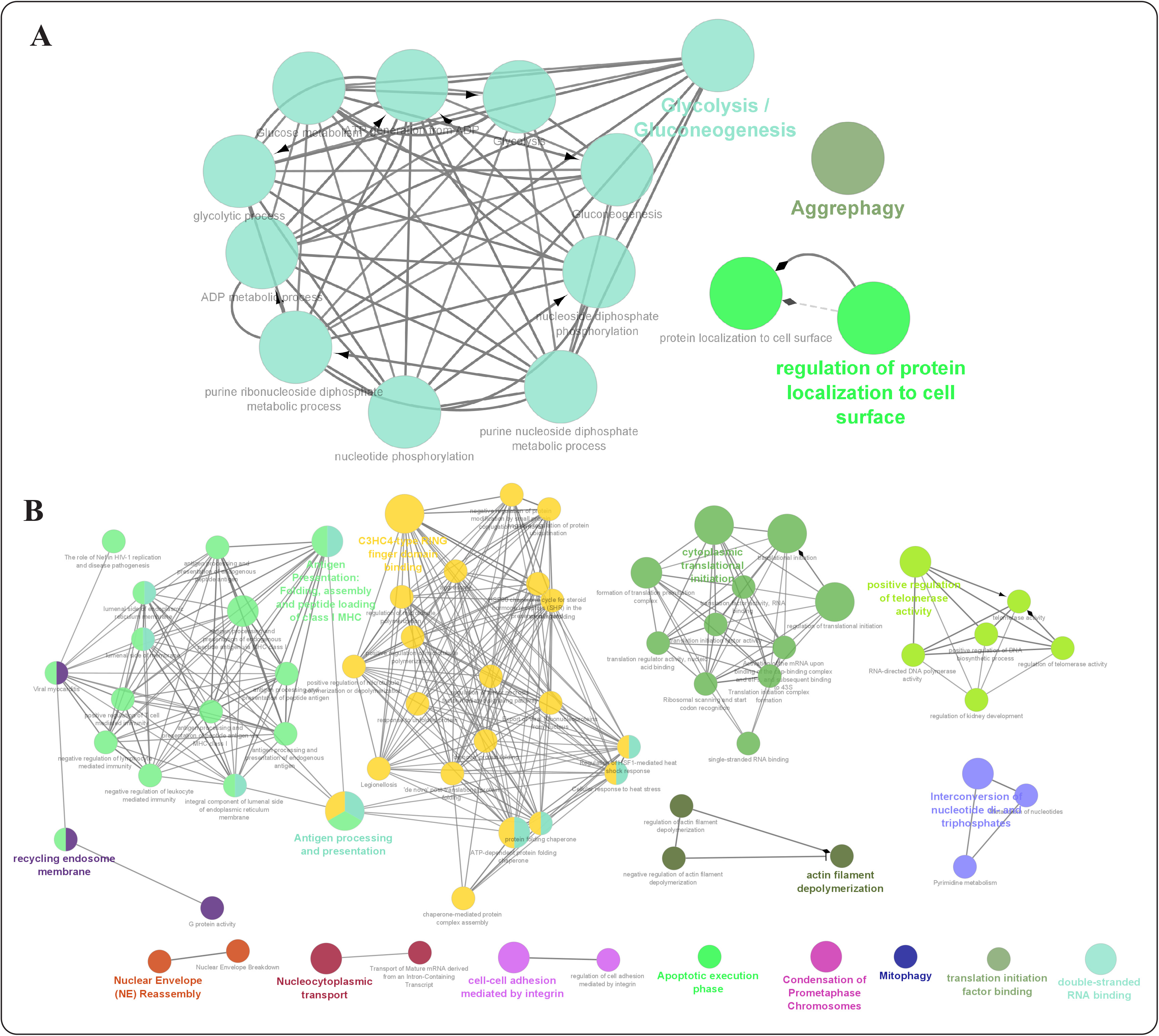
Functional enrichment analysis of most frequently Nm modified genes across a cell line in terms of functional grouping of the GO-terms based on GO hierarchy using Cytoscape ClueGO application. (a) Hek293 cell line and (b) Hela cell line (visualizing high confidence (p-val<0.05) ontologies and pathways potentially associated with Nm RNA modification. The size of the nodes are representative of the significance of association with respect to genes per GO-term.

From the functional enrichment analysis of the top 1% gene set form Hek293 cell line (Figure 5A, and supplementary Figure 4), we observed a wide range of functional processes like: “Glycolysis/Gluconeogenesis”, “Regulation of protein localization to cell surface”, and “Aggrephagy” being significantly enriched. Essentially highlighting the diverse regulatory role of Nm modifications, from their involvement in metabolic pathways, protein degradation and localization.

In Hela cell line, we observed several high confidences (adjusted p-val < 0.05) enriched ontologies that were more representative of Nm modification role in immune response and cellular processes (Figure 5B and supplementary Figure 5) like: “C3HC4-type RING finger domain binding”, “Antigen processing and presentation (class I MHC)”, and “cytoplasmic translational initiation”.

To observe which cellular pathways were associated with the Nm modifications, we ranked the complete human gene lists from both Hela and Hek293 cell lines based on occurrence of Nm modification locations and performed gene set enrichment analysis (GSEA) [27] using WebGestalt [28]. Across both cell lines we observed that genes associated with metabolic processes, protein binding, and biological regulation were enriched in these ranked lists, reinforcing the association between Nm modification and RNA-protein interaction which was previously observed in literature [29] [30]. The Nm modified gene sets from both cell lines had autoimmune pathways, signaling pathways, and Diabetes pathways in common with a relatively higher enrichment scores (NES>1.2) as seen in supplementary Figure 6. Additionally, we also observed some tissue specific pathways were enriched in case of Hela, like prion disease and Inflammatory bowel disease in Hek293. Those pathways were enriched with high normalized enrichment scores (NES>1.2).

## 3. Discussion

We have noticed that Nm-nano outperforms the existing non-Nanopore tools for Nm sites prediction in the literature [18–20] in terms of accuracy with 1:1 ratio of positive and negative test samples (50.1% for [17], 81.91% for [18], 84.8% for [19], and 99% for Xgboost, the best Nm-nano ML model). However, we found the comparison between Nm-Nano and those tools does not make sense in terms of implementation, but it makes sense in terms of accuracy. This is because those tools were only applied to predict Nm sites in short reads of RNA sequences, while Nm-Nano can predict Nm sites on long reads of RNA sequences. Moreover, those tools were trained and tested on the same dataset, and they were not tested for predicting Nm sites on a combination of two benchmark datasets of RNA sequence data for two different cell lines. As for comparison of the performance of Nm-nano with nanoRMS [22], the existing Nanopore based tool for predicting Nm modifications on direct RNA sequencing data, we found that nanoRMS was only tested on predicting Nm in direct RNA sequences of yeast and it was not tested on predicting the Nm sites in direct RNA sequencing data of human cell lines which are more complex than lower eukaryotes like yeast. Hence, it is not possible to directly compare the accuracy of nanoRMS tool on human cell line data. In addition, nanoRMS predicts Nm sites on individual single reads of direct RNA sequence data, where the single read features are used to train the predictors of nanoRMS. Those features were averaged before Nm prediction, making it not feasible to obtain the contribution of each feature in predicting Nm sites. Moreover, only the accuracy values of predicting the stoichiometry of Nm for each read and no other performance statistics such as precision and recall are reported to evaluate the performance of nanoRMS. Therefore, we can only compare the average of the accuracy values of KNN (the best supervised classifier employed in nanoRMS) to the accuracy of predictors integrated in Nm-nano. Based on this comparison, we conclude that the accuracy of each of the two ML models employed in Nm-nano significantly outperforms the average accuracy of KNN predictor (66.17%), employed in nanoRMS. As for comparison between the implementation of Nm-nano and nano-RMS, we found that nano-RMS relies mainly in its implementation on the base-calling “error” signatures in the Nanopore data as features for detecting the Nm-modification. However, those base-calling errors might not be the same for each type of modified K-mer context resulting from base-calling as not all modified bases can be detected as base-calling errors. So, the generated benchmark dataset used in nanoRMS might be biased. Moreover, with the advances in Nanopore technology, the Nanopore base-callers would become more accurate, and thus the base-calling errors would become lower resulting in decreasing the size of the training dataset that is used in a tool such as nanoRMS, which might lead to deceasing the performance of the ML models deployed in this tool for predicting Nm sites. In other words, relying on the erroneous base-calling of Nanopore RNA sequencing for generating a training data on which ML models applied for detecting Nm modification is a challenge either because not all the modified bases in K-mers contexts resulting from base-calling process would generate base-calling errors which might result in biased training or validation dataset, or because of the high accuracy of base-calling might lead to generating limited-size of training data. So, we believe that detecting RNA modifications based on the base-calling error would be an obsolete approach in the future when the base-callers performance becomes or close to 100% accuracy. Finally, we compared the performance of Nm-nano predictors with the performance of HybirdNm proposed in [23] and we found that the accuracies of HybirdNm for predicting Nm sites were not explicitly mentioned. However, we found that AUC was a common metric used to evaluate the performance of Nm-nano and HybirdNm and so we used this metric for comparing the performance of Nm-nano predictor models to HybirdNm performance. In view of this, we found that the best AUC achieved by HybirdNm was 0.962 for predicting Um subtype and the average AUC for predicting the four subtypes was 0.917 which is less than the AUC achieved by either Xgboost (AUC of 0.991) or RF with kmer embedding (AUC of 0.957) employed in Nm-nano framework when any of both models applied to Hek293 benchmark dataset with random test splitting (supplementary file 1_test_split_Hek293_results. docx). Additionally, HybirdNm framework uses the basecalling errors in Nanopore data as a feature for predicting Nm subtypes which can’t be considered a good choice of feature for training the model once the performance of the current Nanopore basecallers reach the optimum by achieving 100% accuracy or very closed to that. Moreover, HybirdNm was trained and tested on the same benchmark dataset of Hek29 cell line, and it was not tested for predicting Nm sites on a combination of two benchmark datasets of RNA sequence data for two different cell lines. Supplementary file (Nm-nano_advantages. docx) tabulates the advantages of Nm-nano over the already available ONT methods for predicting Nm modifications.

It has been also shown that Nm-nano predictors exhibited a high accuracy in the random test-split applied to either Hela or Hek293 benchmark datasets, and the integrated validation test applied to a combination of both Hela and Hek293 bechmark datasets. However, we found that the performance of Nm-nano predictors became significantly lower in the validation with an independent cell line, when once cell line is used for training the Nm-Nano predictor and another cell line is used for testing it. For instance, RF and XGboost achieved an accuracy of 66% and 59% respectively in detecting Nm-sites on Hek293 benchmark dataset in the testing phase after using Hela benchmark dataset for training the models. In the inverse cross validation, RF and XGboost achieved an accuracy of 57.26% and 56% respectively in detecting Nm-sites on Hela benchmark dataset in the testing phase after using Hek293 benchmark dataset for training the models. We believe that this clear decrease in cross validation and inverse cross validation accuracies of detecting Nm sites is due to the small dataset size of Nm-seq data [17] used for generating the benchmark training dataset of Hela and Hek293. This small dataset size causes an increase in the specific differences between both cell lines which causes the decrease in Nm prediction accuracy when tested on independent cell line. In addition, it is also possible that there are cell line specific features which are not captured when trained on individual cell line datasets, resulting in lower cross validation accuracies. Clearly this was not the case in the integrated validation testing of Nm-nano predictors that achieved a high accuracy of detecting Nm sites when training these predictors on 50% of the integrated dataset that combines both benchmark datasets of Hela and Hek293 and test the predictors on the remaining 50%.

It was also observed that deploying Nm-Nano on direct RNA sequencing data of Hela and Hek293 cell lines leads to identifying top frequently modified Nm genes that are associated with a wide range of biological processes in both cell lines. However, it might be unclear how the enrichment of specific functional families for only two considered human cell lines (Hela and Hek293) would strengthen the confidence in the Nm-nano’s predictions. To address this point, we have now looked at publicly available direct RNA sequencing data for human cell lines on SRA and currently we found that there is data for multiple cell lines. However, the data is only available in the form of fastq files and not in the form of fast5 files and given that our algorithm needs signal level fast5 data, it is not possible at this point to run our algorithm at those datasets. Hence, we can’t test on additional cell lines available in the public domain due to the limitations in the access to the fast5 files. However, studying the differential extense of Nm sites across multiple cell lines for understanding common and unique sites is an exciting question which we believe can be addressed as more cell line based direct RNA sequencing datasets in the form of fast5 files with signal data are publicly available in the future. Finally, it is worth noting that the current study has a limitation of only detecting Nm modification in mRNA. This is due to that the current protocol of Nanopore direct RNA sequencing is limited to sequencing mRNA with polyA [31]. However, for other small RNAs to be captured by Nanopore sequencing, it is possible to attach polyA with a modified protocol of Nanopore RNA sequencing. Hence mapping modifications on such small RNAs is beyond the scope of the current study which focuses on mapping Nm modifications on only mRNA. However, we think that it would be possible to apply Nm-Nano predictors for detecting Nm sites in other small RNAs when a modified and rigorously validated version of Nanopore RNA sequencing protocol would be available, that can attach other types of small RNAs to polyA.

## 4. Materials and Methods

### 4.1. Basic approach pipeline

The complete pipeline of Nm-Nano framework for identifying Nm modifications in RNA sequence consists of several stages. The first stage of the pipeline starts by culturing the cell line by extracting it from an animal and letting it grow in an artificial environment. Next, the RNA is extracted from this cell during library preparation and put through the ONT device and starts generating Nanopore signal data. The MinION Mk1B device was used for direct RNA sequencing of Hela and Hek293 cell lines with FLO-MIN**106** flow cell. The fast5 files that store the raw electrical signals output by the ONT device for each cell line are then base-called via Guppy [32] to produce fastq files that store the base-called RNA sequence reads. Those reads are aligned to a reference genome to produce the SAM file using minimap2 tool [33]. From the SAM file, a BAM and sorted BAM file are generated using samtools [34], where the BAM file is a compressed version of the SAM file. Next, a coordinate file is created using the produced SAM file and a provided BED file [35] that includes the Nm-modified locations on the whole genome that have been experimentally verified in literature based on the research work presented in [17]. This coordinate file is needed for labeling the Nanopore signal samples produced by eventalign module as modified or unmodified when training any of the two Nm-Nano predictors (the Supplementary Files 2_Hela.txt and 3_Hek293.txt show the coordinate files generated for Hela and Hek293 respectively). Next, the eventalign module of the Nanopolish (a free software for Nanopore signal extraction and analysis [36–38]) is launched for extracting Nanopore signals, which produces a dataset of Nanopore signal samples. The structure of Nm-Nano’s pipeline is similar to the pipelines used by other RNA modification prediction tools [25]. However, Nm-Nano’s pipeline is different in three phases (Figure 1.A and B): the benchmark dataset generation, the feature extraction, and ML models construction phases. The benchmark dataset generation phase in Nm-Nano’s pipeline is different because Nm modifications can occur at any RNA base, and thus all the samples that are generated from signal extraction process are used to identify the Nm sites using the information in the coordinate file, where some of those samples will be labeled as modified with Nm sites, while the remaining are control samples that will be labeled as unmodified. Similarly, the feature extraction phase in Nm-Nano’s pipeline is different because it uses different features (e.g., position, signal/event_mean, signal/event_stdv, model_mean, model_stdv, kmer_match, mean_diff, and word2vec embedding features of K-mers) extracted from the modified and unmodified signal samples to train the constructed ML models for predicting Nm sites. Finally, the ML models construction phase in Nm-Nano’s pipeline is different because it deploys two different ML models (the XGBoost with tuned parameters and Random Forest with K-mer embedding) for predicting Nm sites in long RNA sequence reads. In the next subsection we will highlight those differences by introducing more details about the benchmark dataset generation, feature extraction and ML model constructions.

### 4.2. Benchmark datasets generation

Two different benchmark datasets were generated for Hela and Hek293 cell lines (Supplementary Tables 5_training_hela.xlsx and 6_training_hek.xlsx). Both datasets were generated by considering all nanopore signals samples generated by passing the long RNA sequence of either Hela or Hek293 through the Oxford Nanopore Technologies (ONT) device. Those signals are extracted using the Nanopolish eventalign module. Each dataset is initially labeled with Nm-sites using the bed file that includes the Nm-modified locations on the whole genome that have been experimentally verified in literature based on Nm-seq protocol [35]. Nm-seq reported about Nm sites from two different cell lines with a total number of 699 Nm sites in Hela and 2102 Nm sites in Hek293. Thus, to label each sample as Nm modified or not, all the samples generated from signal extraction were used as the target samples for identifying Nm modification since Nm modifications can occur at any RNA base. Next, the intersection between their position column on the reference genome and the position in the coordinate file (generated from Nm BED file and SAM file for each cell line) is determined. This intersection will represent the positive samples, while the remaining samples will be the negative samples. In the end we have 52,582 samples: 26,291 are positive and 26,291 are negative ones (after sampling the negative samples which are very huge in comparison with positive ones) for Hek293. Similarly, we got 167,374 samples: 83687 are positive samples and 83,687 are negative samples for Hela cell line. We observed that there is a total of 507 and 1024 different reference kmer combinations captured in the modified and unmodified signals datasets respectively in Hela training data (Supplementary Tables 7_training_modified_kmer_freq_hela.xlsx and 8_ training_unmodified_kmer_freq_hela.xlsx) and a total of 238 and 1022 different reference K-mer combinations captured in the modified and unmodified signal datasets respectively in HeK293 training data (Supplementary Tables 9_training_modified_kmer_freq_hek.xlsx and 10_training_unmodified_kmer_freq_hek.xlsx). Supplementary Figures 7_top_10-modified_bases_training_hela and 8_top_10-modified_bases_training_hek provide the sequence logo for the top ten modified bases corresponding to Nm prediction in the benchmark training datasets of Hela and Hek293 cell lines respectively.

### 4.3. Feature extraction

Each generated benchmark dataset has seven columns that represent the seven features that were used for training the ML models that we developed and integrated in Nm-Nano framework. Those features are position, event_level_mean, event_stdv, model_mean, model_stdv, mean_diff, and K-mer_match. The first five features were directly extracted by picking their columns from the eventalign’s output (Supplementary File 4.txt) (namely: position, event_level_mean, event_stdv, model_mean, and model_stdv columns). The sixth feature is generated by calculating the difference between the mean of the signal (event_level_mean) and the mean of the simulated signal by eventalign module (model_mean). The seventh feature is generated by checking if the reference_K-mer and model_K-mer columns in the eventalign’s output match each other, where the former refers to the base-called K-mers resulting from inferring the RNA sequence reads from the extracted Nanopore signals in the base-calling process, while the latter refers to base-called K-mers resulting from inferring RNA sequence reads from the simulated signals by eventalign. The value of reference & model K-mer match is 1 if reference and model K-mers match each other and 0 otherwise. We should also mention that the position feature simply refers to genomic location of Nm modification and it does not include any information about the nature of nucleotide or neighboring sequence, so, training of Nm-nano predictors with such a feature will not cause the predictions to be highly biased towards the same conserved sequence in other RNA.

### 4.4. Features generation with word embedding

In addition to the extracted features, embedding features have been generated by applying the word2vec technique [39] to the corpus of reference K-mers resulting from aligning Nanopore signals to a reference genome using eventalign module of Nanopolish software. Applying the word2vec technique to the corpus of reference K-mers generates/outputs a set of 1-dimensional vectors of fixed size that represent the embedding features of those reference K-mers (the vector size is set optionally as a parameter when building word2vec embedding model).

The idea of applying Word2vec to reference K-mer has been inspired by the research work in [40], in which word2vec has been applied to DNA K-mers to generate embedding features represented by vectors of real numbers as representations of those K-mers. This approach was introduced as an alternative approach to vector encoding of K-mer using one-hot technique that is subject to the curse of dimensionality problem, as when increasing the length of RNA sequence, the binary feature representation by one-hot encoding grows exponentially resulting in adding too many features to the dataset [41].

The embedding features generated by word2vec are combined with the other extracted features introduced in section previous section for training the RF classifier model that has been developed for predicting Nm sites in long RNA sequence reads. In other words, the combination of all extracted features and embedding features are used to train the RF model, which in turn will be able to predict whether the signal is modified by the presence of Nm sites in the testing phase.

### 4.5 ML Models construction

We have developed two machine learning models for predicting Nm sites in RNA sequence reads including the XGBoost [42] with tuned parameters and RF [43] with K-mer embedding. The XGBoost model parameters were tuned using the Grid-search hyperparameter tuning algorithm [24]. For the RF, the seed number parameter was set to 1234 and the number of trees parameter was set to 30 for obtaining the best performance of RF. The optimized distributed gradient boosting python library has been used for implementing the XGBoost model [44] and the scikit-learn toolkit [45], the free machine learning python library has been used for implementing the RF model.

#### 4.5.1. XGBoost with grid search for hyper parameter tuning

The Extreme Gradient Boosted trees (XGBoost) is a special implementation of Gradient Boosting [46]. Gradient boosting is a machine learning technique that produces a prediction model based on an ensemble of weak prediction models, which are decision trees in the case of XGBoost. This model is highly flexible and versatile and can be applied for classification-based problems, which is the main goal of this study. The advantage that XGBoost has over other tree-based models is that it has a faster training time along with its regularized boosting, which helps to prevent overfitting: this is when the machine learning model learns and becomes too accustomed to the training data and is not able to generalize and accurately predict the testing data. XGBoost does not also require feature scaling due to being a tree-based model and so feature scaling did not affect the value of the split point and the structure of the tree model. XGBoost can also cross-validate each iteration (round) of its training process, which can lead to higher results than models that cannot do the latter process. The use of decision trees and gradient boosting also provide the advantages over both random forest and other gradient boosting models, causing XGBoost to typically have a prediction error many times lower than regular gradient boosting or random forest.

The XGBoost machine learning model was created after the data was preprocessed by removing all null values and performing feature extraction, The model has several parameters that can be adjusted and tuned to get the best performance of XGBoost. Hyper-parameter tuning using the grid search algorithm has been used since it allows for the best and most accurate combination of parameters to be obtained. The parameters that were optimized for the XGBoost model were eta, gamma, max_depths, min_child_weights, and scale_pos_weight. The optimized values for these parameters obtained using grid search algorithm were 0.01, 0.1, 15, 3, and 1 respectively. The parameter eta, representing the learning rate of the XGBoost model. Gamma parameter represents how conservative the model is. The parameter max_depth represents how deep a decision tree can be built and min_child_weight represents the minimum value needed to activate the respective node in the decision tree. The scale_pos_weight parameter controls the balance of positive and negative weights; this parameter is associated with the min_child weight. After the values for these best parameters were obtained by fitting the grid search XGBoost model to the training data, they were applied to the model to obtain its prediction results in the testing phase.

XGBoost is trained with the set of features mentioned in section 4.3. Those features are extracted from the raw signal provided by direct RNA nanopore sequencing and the corresponding base-called K-mers resulting from inferring the underlying RNA sequence by base-calling.

#### 4.5.2. RF with K-mer embedding

We have developed a Random Forest (RF) ML model that has been trained with the same set of features used to train the XGBoost model, in addition to the embedding features generated by applying Word2vec embedding technique to the reference K-mers of in the extracted Nanopore signals., one feature column in the benchmark Nm modification datasets of Hela and Hek293 cell lines. RF algorithm has been extensively used in the literature to address several problems in bioinformatics research [47]. It has been observed that the features generated by applying Word2vec embedding technique to the reference K-mers have a great positive impact on the performance of RF model as it was mentioned in results subsections 2.1 and 2.2.

The RF ML model was created after the data was preprocessed by removing all null values then performing feature extraction and combining them with generated K-mer embedding features. The K-mer embedding features were generated using genism [48], a free python library that implements word2vec algorithm using highly optimized C routines, data streaming, and pythonic interfaces. The word2vec algorithm has various parameters including: the vector size, the window size, and the word count. The vector size is the dimensionality of the vector that represents each K-mer. The window size refers to the maximum distance between a target word/K-mer and words/K-mers around the target word/K-mer. The word count refers to the minimum count of words to consider when training the model, where words with occurrence less than this count will be ignored. The K-mer embedding features that lead to best performance of RF have been generated by setting the vector size to 20, the minimum word count to 1, and the window size to 3.

### 4.6. Performance evaluation metrics

The accuracy (Acc), precision (P), recall (R), and the area under ROC curve (AUC) [49] have been used as metrics for evaluating performance of Nm-Nano predictors. The mathematical notions for the first three metrics are identified as follows:

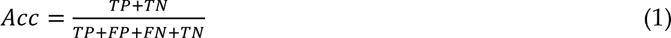

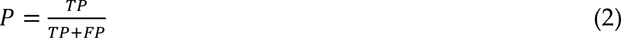

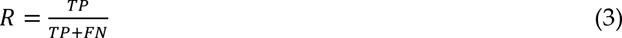

Where:

- TP denotes true positive and refers to the number of correctly classified Nm sites.
- FP denotes false positive and refers to the number of non-Nm sites misclassified as Nm sites.
- FN denotes false negative and refers to the number of Nm sites misclassified as non-Nm sites.
- TN denotes true negative and refers to the number of correctly classified non-Nm sites. As for AUC metric, it measures the entire two-dimensional area under the ROC curve [50] which measures how accurately the model can distinguish between two things (e.g. determine if a base of RNA sequence is Nm site or not).

### 4.7. Environmental settings

Nm-Nano has been developed as tool for detecting Nm modification in Nanopore RNA sequence data by integrating two ML models: the XGBoost with tuned parameters and RF with K-mer embedding to predict this type of RNA modification. XGBoost parameters were tuned to get the best performance using the Grid search algorithm which takes around 6 hours and 52 minutes to fit on the training dataset of Hek293 and 9 hours and 12 minutes to fit on the training dataset of Hela cell line for obtaining the best parameters that were applied to XGBoost model in the testing phase. The experiment was executed on Windows 10 machine with (8 cores) processor of Ryzen 5900HS CPU, and 16 GB RAM. It should be mentioned that though grid search algorithm took a huge processing time for tuning the parameters of XGboost, using this algorithm for optimizing XGBoost parameter causes a significant improvement in XGBoost performance. The performance results of XGBoost versus XGBoost tuned with grid search algorithm are shown in Supplementary file 5_xgboost_versus_grid_search_xgboost_results.docx. Similarly, though using word2vec for generating embedding features when developing RF with K-mer embedding model added extra processing time to the execution time of RF algorithm, it also led a great improvement in RF performance. This is because combining the embedding features generated by word2vec with the extracted features from Nanopore signals positively affects the performance of RF since the embedding features are strongly contributes to the model performance as it has been presented in result subsections 2.1 and 2.2. The performance results of RF versus RF with kmer embedding are shown in Supplementary file 6_RF_versus_RF_with_kmer_embedding_results.docx. Meanwhile, we thought about improving the performance of XGBoost by applying grid search algorithm for hyper parameter tuning in addition to applying K-mer embedding with word2vec for generating embedding features that would be combined with the extracted features used for training XGBoost. However, we found that this will make XGBoost very slow when applying it to the benchmark dataset of a given cell line with a slight improvement in its performance that would not be proportional to the huge increase in the processing time of XGBoost as it has been presented in result subsection 2.1.

### 4.8. Implementation and usage of Nm-Nano

The ML models of Nm-Nano framework are implemented in python 3.x. To run the XBboost model, the user has to type the following command from Nm-Nano main directory on the user’s local machine after cloning the code from Nm-Nano GitHub repository:

python test_xgboost.py

Similarly, to run Rf with K-mer embedding model, the user has to type the following command from Nm-Nano main directory:

python RF_embedding.py

To allow user to practice with Nm-nano predictors, we include a small benchmark dataset sample for Hela cell line in Nm-nano GitHub repository (Nm_benchmark_hela_sample.csv). However, the user is free to generate a benchmark dataset for any other cell lines based on the instructions mentioned in README file in generate_benchmark folder in Nm-nano GitHub repository.

To generate a benchmark dataset for a specific cell line, the following command should be run on the command line of Linux environment from generate_benchmark folder in Nm-Nano main directory:

python main.py -r ref.fa -f reads.fastq

Where main.py is a python script file included in generate_benchmark folder implemented in python 3.x, (ref.fa) is the reference Genome file and (reads.fastq) is the fastq reads file. Both ref.fa and reads.fastq files should be placed in the same path with the main.py file.

Before running the main script, the user should include the folder that includes the fast5 files (fast5_files) from which reads.fastq file was generated in the same directory of main.py file. Once the user runs main.py script, it will lunch executing several command lines for generating eventalign output. These command lines are included in generate_eventalign_output.txt in in generate_benchmark folder on Nm-nano GitHub repository. Meanwhile the main.py will call other two python files. The first is gen_coors_Nm.py that generates the coordinate file by asking the user to enter the name of the bed file that has the Nm-modified genomic locations with the absolute path and extension. The second is extract_nm.py that takes as input the coordinate file and the eventalign output to extract features and generate the benchmark dataset. To allow the user to practice with Nm-nano pipeline for benchmark dataset generation, we include the following in generate_benchmark folder on Nm-nano GitHub repository

1. - A link to download a fast5 file sample for the Hek293 cell line that should be included in fast5_files folder that should be placed in the same path of main.py file.
2. - A sample of fastq files (reads.fastq) for Hek293 corresponding to the fast5 files in step1
3. - A link to download a reference genome sample (ref.fa) that should be placed in the same path of main.py file.
4. - A sample of bed file for Hek293 cell line (hek.bed.txt)

It should also be mentioned that Nm-nano framework can also be extended by integrating other ML/ deep learning models for predicting Nm sites. Moreover, the framework’s pipeline is generic and can be used with any direct RNA sequencing output from any ONT device such as MinION, GridION, and PromethION.

## 5. Conclusions

In this paper, we have proposed a new framework called Nm-Nano that integrates two machine learning models: the XGBoost with tuned parameters using grid search algorithm and RF with K-mer embedding. It has been shown that the proposed framework was efficient in detecting Nm sites in RNA long reads of human cell lines which addresses the limitations of existing Nm predictors presented in the literature that were only able to detect Nm sites in short reads of RNA sequences of cell lines of various species or long reads of RNA sequences of non-human cell lines (yeast) and one human cell line (Hek293).

By deploying Nm-Nano on direct RNA sequencing data of Hela and Hek293 cell lines, the top frequently modified Nm genes that are associated with a wide range of biological processes in both cell lines were identified. In Hela, we observed several high confidences (adjusted p-val < 0.05) enriched ontologies that were more representative of Nm modification role in immune response and cellular processes like: “C3HC4-type RING finger domain binding”, “Antigen processing and presentation (class I MHC)”, and “cytoplasmic translational initiation”, while in Hek293 we observed a wide range of functional processes like: “Glycolysis/Gluconeogenesis”, “Regulation of protein localization to cell surface”, and “Aggrephagy” being significantly enriched that highlights the diverse regulatory role of Nm modifications, from their involvement in metabolic pathways, protein degradation and localization. Thus, Nm-Nano would be a useful computational framework for accurate and interpretable predictions of Nm sites in RNA sequence read of human and other species cell lines as well that could reveal various biological findings.

## Author Contributions

DH, AA, and SCJ conceived and designed the study. DH implemented the Nm-Nano Github software version. AA and DH implemented the Nm modifications ML predictors namely XGBoost and RF with K-mer embedding respectively. DH extracted the benchmark datasets. AA tuned the parameters of XGBoost using the Grid-search algorithm. AA and DH evaluated the performance of XGBoost and RF with K-mer embedding models with the random test split and integrated validation testing. DH identified the unique Nm genomic locations and the top modified RNA bases with Nm sites on Hela and Hek293 cell lines. SVD performed results, functional and gene set enrichment analysis. QM performed the cell culturing, RNA library preparation and Nanopore RNA sequence for Hela and Hek293 cell lines.

## Data Availability Statement

Nm-nano is available at the Github repository https://github.com/Janga-Lab/Nm-Nano. The directRNA-sequencing data generated in this study for Hek293, and Hela cell lines is publicly available on SRA, under the project accession PRJNA685783 and PRJNA604314 respectively.

## Acknowledgments

We thank Alexander Krohannon, Hunter M. Gill and Alexandre Plastow at IUPUI for giving valuable comments on this work.

## References

1. Darzacq, X.J., B.E.; Verheggen, C.; Kiss, A.M.; Bertrand, E.; Kiss, T., Cajal body-specific small nuclear RNAs: A novel class of 20-O-methylation and pseudouridylation guide RNAs. EMBOJ. 2002, 21, 2746–2756.

2. Rebane, A.R., H.; Metspalu, A., Locations of several novel 2’-O-methylated nucleotides in human 28S rRNA. BMC Mol. Biol. 2002, 3, 1.

3. Somme J, V.L.B., Roovers M, Steyaert J, Versées W, Droogmans L., Characterization of two homologous 2’-O-methyltransferases showing different specificities for their tRNA substrates. RNA. 2014 Aug;20(8):1257-71. doi: 10.1261/rna.044503.114. Epub 2014 Jun 20. PMID: 24951554; PMCID: PMC4105751.

4. Kurth HM, M.K., 2’-O-methylation stabilizes Piwi-associated small RNAs and ensures DNA elimination in Tetrahymena. RNA. 2009 Apr;15(4):675-85. doi: 10.1261/rna.1455509. Epub 2009 Feb 24. PMID: 19240163; PMCID: PMC2661841.

5. Elliott BA, H.H., Ranganathan SV, Vangaveti S, Ilkayeva O, Abou Assi H, Choi AK, Agris PF, Holley CL., Modification of messenger RNA by 2’-O-methylation regulates gene expression in vivo. Nat Commun. 2019 Jul 30;10(1):3401. doi: 10.1038/s41467-019-11375-7. PMID: 31363086; PMCID: PMC6667457.

6. Guy MP, S.M., Weiner CL, Hobson L, Stark Z, Rose K, Kalscheuer VM, Gecz J, Phizicky EM., Defects in tRNA anticodon loop 2’-O-methylation are implicated in Nonsyndromic X-linked intellectual disability due to mutations in FTSJ1. Hum Mutat. 2015;36(12):1176–87.

7. Picard-Jean F, B.C., Tremblay-Letourneau M, Allaire A, Beaudoin MC, Boudreault S, Duval C, Rainville-Sirois J, Robert F, Pelletier J, et al., 2’-Omethylation of the mRNA cap protects RNAs from decapping and degradation by DXO. PLoS One. 2018;13(3):e0193804.

8. Hengesbach M, S.H., Structural basis for regulation of ribosomal RNA 2’-o-methylation. Angew Chem Int Ed Engl. 2014;53(7):1742–4.

9. Erales J, M.V., Panthu B, Gillot S, Belin S, Ghayad SE, Garcia M, Laforets F, Marcel V, Baudin-Baillieu A, et al., Evidence for rRNA 2’-Omethylation plasticity: control of intrinsic translational capabilities of human ribosomes. Proc Natl Acad Sci U S A. 2017;114(49):12934–9.

10. Dilyana G. Dimitrova, L.T.a.C.C., RNA 2’-O-Methylation (Nm) Modification in Human Diseases. Genes 2019, 10, 117; doi:10.3390/genes10020117.

11. Krogh, N.B., U.; Nielsen, H., RiboMeth-seq: Profiling of 20 -O-Me in RNA. Methods Mol. Biol. 2017, 1562, 189–209.

12. Motorin Y, M.V., Detection and Analysis of RNA Ribose 2’-O-Methylations: Challenges and Solutions. Genes (Basel). 2018 Dec 18;9(12):642. doi: 10.3390/genes9120642. PMID: 30567409; PMCID: PMC6316082.

13. Yinzhou Zhu, S.P.P.a.G.G.C., High-throughput and site-specific identification of 2’-O-methylation sites using ribose oxidation sequencing (RibOxi-seq). RNA 2017 23: 1303-1314 originally published online May 11, 2017.

14. Yuan, B.-F., Liquid chromatography–mass spectrometry for analysis of RNA adenosine methylation. In RNA Methylation: Methods and Protocols (Lusser, A., ed.), pp. 33–42, Springer New York, 2017.

15. Jora M, L.P., Ross RL, Williams B, Addepalli B., Detection of ribonucleoside modifications by liquid chromatography coupled with mass spectrometry. Biochim Biophys Acta Gene Regul Mech. 2019 Mar;1862(3):280-290. doi: 10.1016/j.bbagrm.2018.10.012..

16. Anreiter I, M.Q., Simpson JT, Janga SC, Soller M., New Twists in Detecting mRNA Modification Dynamics. Trends Biotechnol. 2020 Jul 1;S0167-7799(20)30166-9.doi: 10.1016/j.tibtech.2020.06.002.

17. Dai, Q., Moshitch-Moshkovitz, S., Han, D., et al., Correction: Corrigendum: Nm-seq maps 2’-O-methylation sites in human mRNA with base precision. Nat Methods 15, 226–227 (2018). 10.1038/nmeth0318-226c.

18. Chen, W., et al., Identifying 2’-O-methylationation sites by integrating nucleotide chemical properties and nucleotide compositions. Genomics,107(6): p. 255-258, 2016.

19. Milad Mostavi, S.S.a.Y.H., Deep-2’-O-Me: Predicting 2’-O-methylation sites by Convolutional Neural Networks. In proceedings of Annual International Conference of the IEEE Engineering in Medicine and Biology Society, July 2018.

20. Zhou, Y., Cui, Q. & Zhou, Y., NmSEER V2.0: a prediction tool for 2’-O-methylation sites based on random forest and multi-encoding combination. BMC Bioinformatics 20, 690 (2019).

21. Wan YK, H.C., Pratanwanich PN, Göke J., Beyond sequencing: machine learning algorithms extract biology hidden in Nanopore signal data. Trends Genet. 2022 Mar;38(3):246-257. doi: 10.1016/j.tig.2021.09.001.

22. Begik O, L.M., Pryszcz LP, Ramirez JM, Medina R, Milenkovic I, Cruciani S, Liu H, Vieira HGS, Sas-Chen A, Mattick JS, Schwartz S, Novoa EM., Quantitative profiling of pseudouridylation dynamics in native RNAs with nanopore sequencing. Nat Biotechnol. 2021 Oct;39(10):1278-1291. doi: 10.1038/s41587-021-00915-6. Epub 2021 May 13. PMID: 33986546.

23. Pan, S., et al., Prediction and Motif Analysis of 2’-O-methylation Using a Hybrid Deep Learning Model from RNA Primary Sequence and Nanopore Signals. Current Bioinformatics 17.9 (2022): 873-882.

24. Dagnew., B.H.S.a.G., Grid Search-Based Hyperparameter Tuning and Classification of Microarray Cancer Data. In Proceedings of Second International Conference on Advanced Computational and Communication Paradigms (ICACCP), 2019.

25. Hassan D, A.D., Daulatabad SV, Mir Q, Janga SC., Penguin: A tool for predicting pseudouridine sites in direct RNA nanopore sequencing data. Methods. 2022 Jul;203:478-487. doi: 10.1016/j.ymeth.2022.02.005. Epub 2022 Feb 16. PMID: 35182749; PMCID: PMC9232934.

26. Bindea G, M.B., Hackl H, Charoentong P, Tosolini M, Kirilovsky A, Fridman WH, Pagès F, Trajanoski Z, Galon J., ClueGO: a Cytoscape plug-in to decipher functionally grouped gene ontology and pathway annotation networks. Bioinformatics. 2009 Apr 15;25(8):1091-3. doi: 10.1093/bioinformatics/btp101.

27. Subramanian A, T.P., Mootha VK, Mukherjee S, Ebert BL, Gillette MA, Paulovich A, Pomeroy SL, Golub TR, Lander ES, Mesirov JP., Gene set enrichment analysis: a knowledge-based approach for interpreting genome-wide expression profiles. Proc Natl Acad Sci U S A. 2005 Oct 25;102(43):15545-50. doi: 10.1073/pnas.0506580102.

28. Liao Y, W.J., Jaehnig EJ, Shi Z, Zhang B., WebGestalt 2019: gene set analysis toolkit with revamped UIs and APIs. Nucleic Acids Res. 2019 Jul 2;47(W1):W199-W205. doi: 10.1093/nar/gkz401. PMID: 31114916; PMCID: PMC6602449.

29. Hou YM, Z.X., Holland JA, Davis DR., An important 2’-OH group for an RNA-protein interaction. Nucleic Acids Res. 2001 Feb 15;29(4):976-85. doi: 10.1093/nar/29.4.976. PMID: 11160931; PMCID: PMC29614.

30. Lacoux C, D.M.D., Boyl PP, Zalfa F, Yan B, Ciotti MT, Falconi M, Urlaub H, Achsel T, Mougin A, Caizergues-Ferrer M, Bagni C., BC1-FMRP interaction is modulated by 2’-O-methylation: RNA-binding activity of the tudor domain and translational regulation at synapses. Nucleic Acids Res. 2012 May;40(9):4086-96. doi: 10.1093/nar/gkr1254. Epub 2012 Jan 11. PMID: 22238374; PMCID: PMC3351191.

31. Garalde, D., Snell, E., Jachimowicz, D. et al., Highly parallel direct RNA sequencing on an array of nanopores. Nat Methods 15, 201–206 (2018). 10.1038/nmeth.4577.

32. Basecalling using Guppy. Workflows and tutorials for LongRead analysis with specific focus on Oxford Nanopore data. Available from: https://timkahlke.github.io/LongRead_tutorials/BS_G.html.

33. H., L., Minimap2: pairwise alignment for nucleotide sequences. Bioinformatics. 2018 Sep 15;34(18):3094-3100. doi: 10.1093/bioinformatics/bty191.

34. Danecek P, B.J., Liddle J, Marshall J, Ohan V, Pollard MO, Whitwham A, Keane T, McCarthy SA, Davies RM, Li H, Twelve years of SAMtools and BCFtools, GigaScience (2021) 10(2).

35. BED file format - Genome Browser FAQ. Available from: https://genome.ucsc.edu/FAQ/FAQformat.html#format1.

36. Loman, N., Quick, J. & Simpson, J., A complete bacterial genome assembled de novo using only nanopore sequencing data. Nat Methods 12, 733–735 (2015). 10.1038/nmeth.3444.

37. Simpson., J., Aligning Nanopore Events to a Reference. Apr 8, 2015.

38. Nanopolish. Available from: https://github.com/jts/nanopolish.

39. T. Mikolov, K.C., G. Corrado, and J. Dean., Efficient estimation of word representations in vector space. (2013a). arXiv preprint arXiv:1301.3781.

40. Ng., P., dna2vec-Consistent vector representations of variable-length k-mers. 10.48550/arXiv.1701.06279.

41. Milad Mostavi, Y.H., Machine Learning and Deep Learning challenges for building 2’O site prediction. bioRxiv 2020.05.10.087189; doi: 10.1101/2020.05.10.087189.

42. Guestrin., T.C.a.C., XGBoost: A Scalable Tree Boosting System. In Proceedings of the 22nd ACM SIGKDD International Conference on Knowledge Discovery and Data Mining (KDD ’16), August 13-17, 2016, San Francisco, CA, USA.

43. Breiman, L., Random Forests. Machine Learning, 45, 5-32, 2001.

44. Jain., A., in Complete Guide to Parameter Tuning in XGBoost with codes in Python. March 2016. Avialble at: https://www.analyticsvidhya.com/blog/2016/03/complete-guide-parameter-tuning-XGBoost-with-codes-python/.

45. scikit-learn Machine Learning in Python. Available from: https://scikit-learn.org/stable/.

46. Grover, P., Gradient Boosting from scratch. Dec 8, 2017.

47. Qi, Y., Random Forest for Bioinformatics. In Ensemble Machine Learning, pp. 307-323, Springer, 2012.

48. Genism topic modelling for humans. Available from: https://radimrehurek.com/gensim/models/word2vec.html.

49. Bradley., A.E., The Use of the Area under the Roc Curve in the Evaluation of Machine Learning Algorithms. Pattern Recognition, Vol. 30, No. 7, pp. 1145-1159, 1997.

50. Receiver operating characteristic. Available at: https://en.wikipedia.org/wiki/Receiver_operating_characteristic

