## Supplementary material for "Nm-Nano: A Machine Learning Framework for Transcriptome-Wide Single Molecule Mapping of 2´-O-Methylation (Nm) Sites in Nanopore Direct RNA Sequencing Datasets": Supplementary_Figures_Legends.docx

**Supplementary Figures legend**

| **1_test_split_hek** | The learning, loss and ROC curves of XGBoost (Panel A, B, and C) against the learning, loss and ROC curves of RF with embedding (Panel D, E, and F) with random test split on Hek293 benchmark dataset. |
| --- | --- |
| **2_top_10 modified_bases_hela** | The sequence logo for the top ten modified bases corresponding to Nm prediction in Hela. |
| **3_top_10-modified_bases_hek** | The sequence logo for the top ten modified bases corresponding to Nm prediction in Hek203. |
| **Figure 4** | The functional enrichment analysis of the gene set contributing to Nm prediction in Hek293 cell line. |
| **Figure 5** | The functional enrichment analysis of the gene set contributing to Nm prediction in Hela cell line. |
| **Figure 6** | The gene set enrichment analysis (GSEA) of Nm modified genes set across cell line (a) Hek293 (b) Hela cell |
| **7_top_10-modified_bases_training_hela** | The sequence logo for the top ten modified bases corresponding to Nm prediction in the benchmark training dataset of Hela cell line. |
| **8_top_10-modified_bases_training_hek** | The sequence logo for the top ten modified bases corresponding to Nm prediction in the benchmark training datasets of Hek293 cell line. |
