## Supplementary material for "Nm-Nano: A Machine Learning Framework for Transcriptome-Wide Single Molecule Mapping of 2´-O-Methylation (Nm) Sites in Nanopore Direct RNA Sequencing Datasets": 1_test_split_Hek293_results.docx

**Supplementary Table 1.** The performance of Nm-nano predictors on Hek293 benchmark dataset with random-test splitting.

| **Classifier** | **Accuracy** | **Precision** | **Recall** | **AUC** |
| --- | --- | --- | --- | --- |
| XGBoost | 99 | 0.99 | 0.99 | 0.991 |
| RF | 96 | 0.94 | 0.98 | 0.957 |
