## Supplementary material for "Nm-Nano: A Machine Learning Framework for Transcriptome-Wide Single Molecule Mapping of 2´-O-Methylation (Nm) Sites in Nanopore Direct RNA Sequencing Datasets": 5_xgboost_versus_grid_search_xgboost_results.docx

| **AUC** | **recall** | **precision** | **accuracy (%)** | | **Classifier** | |
| --- | --- | --- | --- | --- | --- | --- |
| 0.795 | 0.72 | 0.85 | | 79.49 | | XGBoost |
| 0.986 | 0.98 | 0.99 | | 98.6 | | XGBoost with grid search |

**Supplementary Table 2**: The performance of XGBoost versus the performance of the XGBoost with grid search algorithm applied to Hela benchmark dataset with random test-splitting.

**Supplementary Table 3**: The performance of XGBoost versus the performance of the XGBoost with grid search algorithm when tested on independent/different cell line.

| **AUC** | **recall** | **precision** | **accuracy (%)** | **Classifier** | |
| --- | --- | --- | --- | --- | --- |
| 0.78 | 0.69 | 0.84 | 78.05 | | XGBoost |
| 0.932 | 0.9 | 0.96 | 93.2% | | XGBoost with grid search |
