## Supplementary material for "Nm-Nano: A Machine Learning Framework for Transcriptome-Wide Single Molecule Mapping of 2´-O-Methylation (Nm) Sites in Nanopore Direct RNA Sequencing Datasets": 6_RF_versus_RF_with_kmer_embedding_results.docx

**Supplementary Table 4.** The performance of RF versus the performance of the RF with k-mer embedding model applied to Hela benchmark dataset with random test-splitting.

| **AUC** | **recall** | **precision** | A **p Accuracy (%)** | **Classifier** |
| --- | --- | --- | --- | --- |
| 0.89 | 0.93 | 0.85 | 88.34 | RF |
| 0.92 | 0.96 | 0.9 | 92.39 | RFwith k-mer embedding |
