## Supplementary material for "Nm-Nano: A Machine Learning Framework for Transcriptome-Wide Single Molecule Mapping of 2´-O-Methylation (Nm) Sites in Nanopore Direct RNA Sequencing Datasets": Nm-nano_advantages.docx

**Supplementary Table 5.** Advantages of Nm-nano over the already available ONT methods for Nm predictions

| Software | Pure ONT framework | Species on which the software can be applied | Predict Nm-sites based on base-calling errors | Report about feature importance |
| --- | --- | --- | --- | --- |
| nanoRMS | Yes | Yeast | Yes | No |
| HybridNm | No | Hek293 | Yes | Yes, using the gradients of  the output with respect to the input sequence |
| Nm-nano | Yes | Human and other species | No, it predicts Nm sites based on features extracted from nanopore signal, making it broad purpose Nm-modification predictors | Yes |
