## Supplementary material for "Nm-Nano: A Machine Learning Framework for Transcriptome-Wide Single Molecule Mapping of 2´-O-Methylation (Nm) Sites in Nanopore Direct RNA Sequencing Datasets": Supplementary_Files_Legends.docx

| **1_test_split_Hek293_results.docx** | The performance of Nm-nano predictors on HK293 benchmark dataset with random-test splitting. |
| --- | --- |
| **2_Hela.txt** | The coordinate file generated for Hela cell line based on the Nm-seq data |
| **3_Hek293.txt** | The coordinate file generated for HeK293 cell line based on Nm-seq data |
| **4.txt**  **5_xgboost_versus_grid_search_xgboost_results.docx** | The eventalign’s output  The performance results of XGBoost versus XGboost tuned with grid search algorithm |
| **6_RF_versus_RF_with_kmer_embedding_results.docx**  **Nm-nano_advantages. docx** | The performance results of RF versus RF with kmer embedding  The advantages of Nm-nano over the already available ONT methods for predicting Nm modifications |
