## Supplementary material for "Nm-Nano: A Machine Learning Framework for Transcriptome-Wide Single Molecule Mapping of 2´-O-Methylation (Nm) Sites in Nanopore Direct RNA Sequencing Datasets": Supplementary_Tables_Legends.docx

| **1_Nm_unique_genomic_locations _hela.xlsx** | The unique genomic location of Nm sites in the complete RNA sequence of Hela cell line. |
| --- | --- |
| **2_Nm_unique_genomic_locations _hek.xlsx** | The unique genomic location of Nm sites in the complete RNA sequence of Hek293 cell line |
| **3_Nm_unique_reference_kmer_freq_hela.xlsx** | The frequency of unique reference_kmers that contribute to Nm predictions in Hela cell line. |
| **4_Nm_unique_reference_kmer_freq_hek.xls** | The frequency of unique reference_kmers that contribute to Nm predictions in Hek293 cell line. |
| **5_training_hela.xlsx** | The benchmark datasets generated for Hela. |
| **6_training_hek.xlsx** | The benchmark datasets generated for Hek293.  . |
| **7_training_modified_kmer_freq_hela.xlsx** | The different reference kmer combinations captured in the modified datasets of Hela cell line |
| **8_ training_unmodified_kmer_freq_hela.xlsx** | The different reference kmer combinations captured in the unmodified dataset of Hela cell line. |
| **9_training_modified_kmer_freq_hek.xlsx** | The different reference kmer combinations captured in the modified datasets of Hek293 cell line. |
| **10_training_unmodified_kmer_freq_hek.xlsx** | The different reference kmer combinations captured in the unmodified dataset of Hek293 cell line. |
