## Supplementary figures and images for "Nm-Nano: A Machine Learning Framework for Transcriptome-Wide Single Molecule Mapping of 2´-O-Methylation (Nm) Sites in Nanopore Direct RNA Sequencing Datasets"

### 1_test_split_hek.png

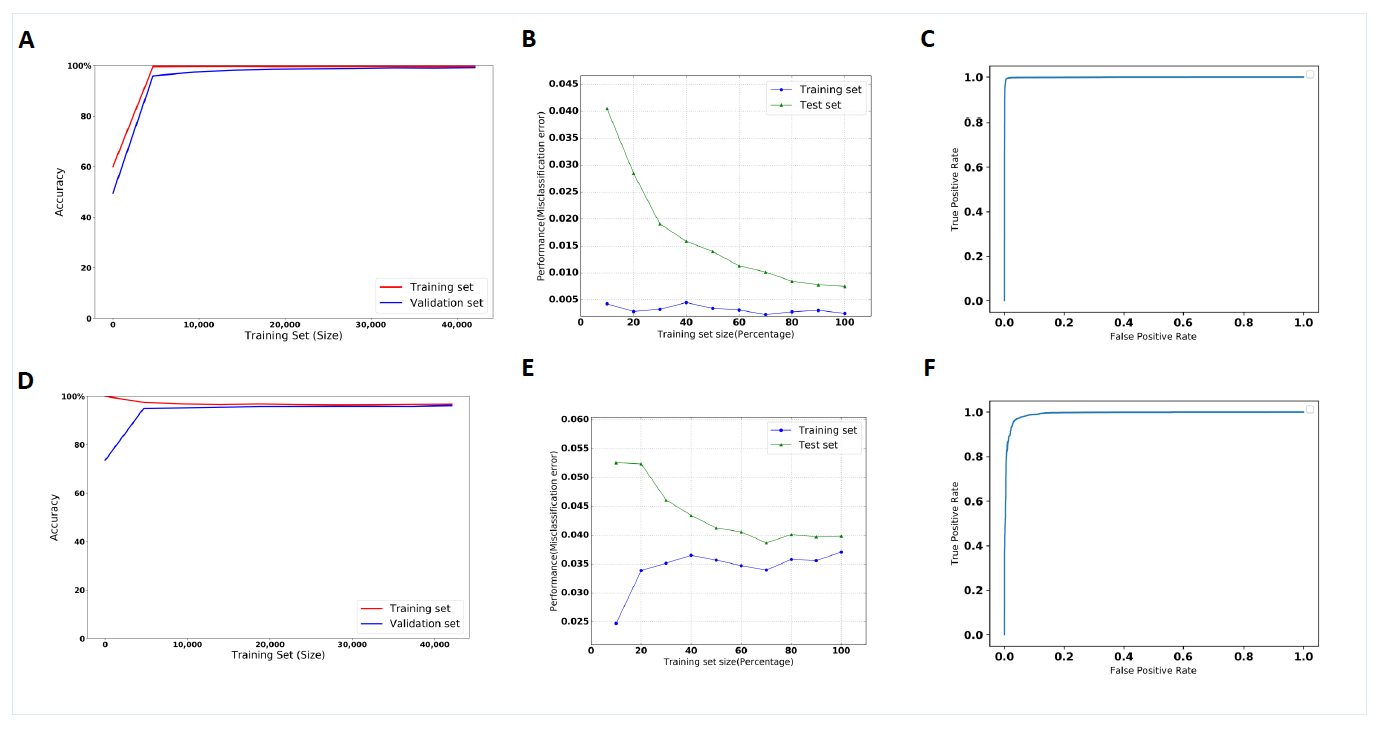

### 2_top_10-modified_bases_hela.png

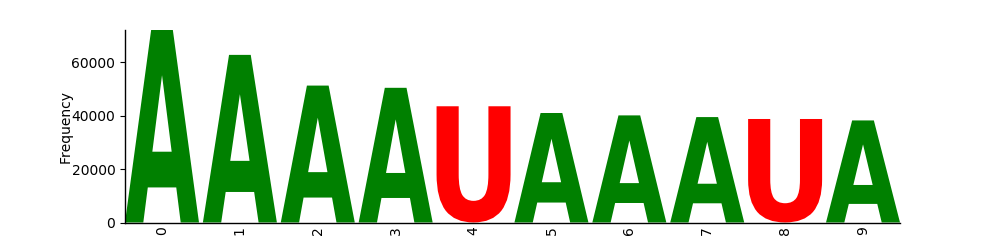

### 3_top_10-modified_bases_hek.png

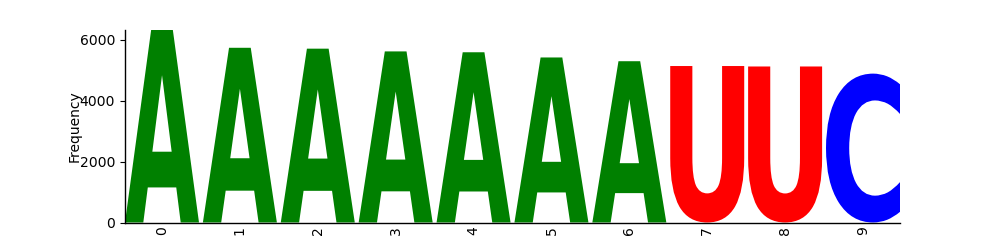

### 4.png

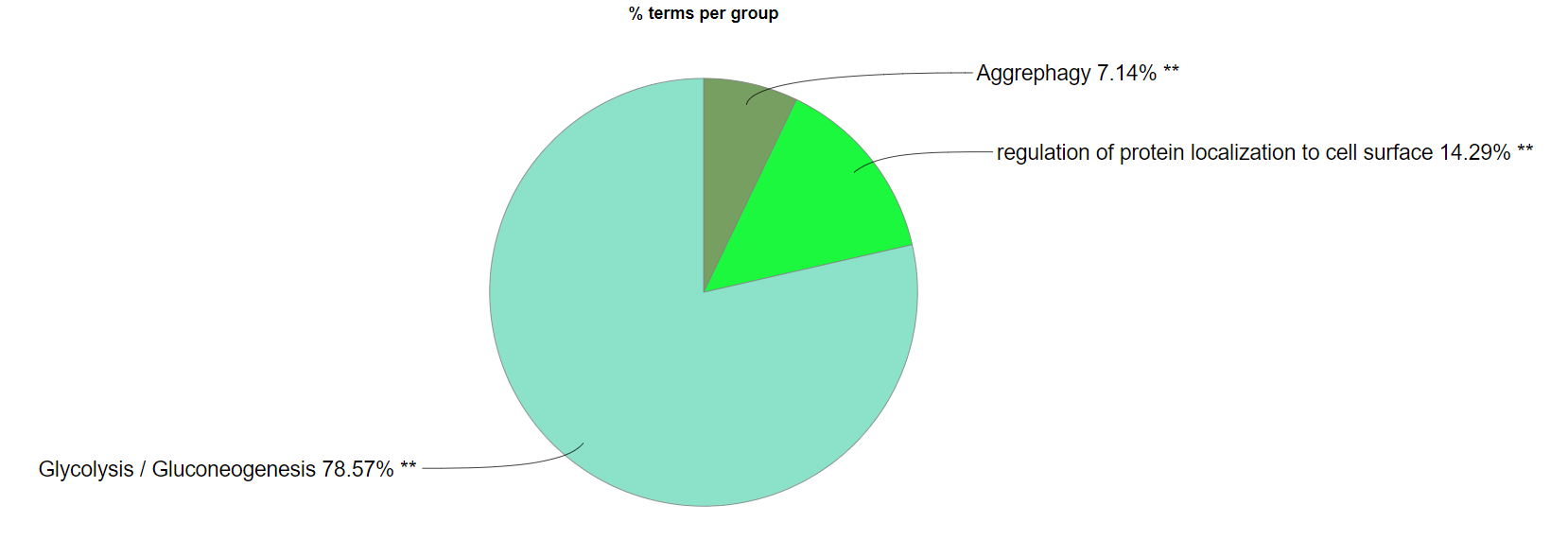

### 5.png

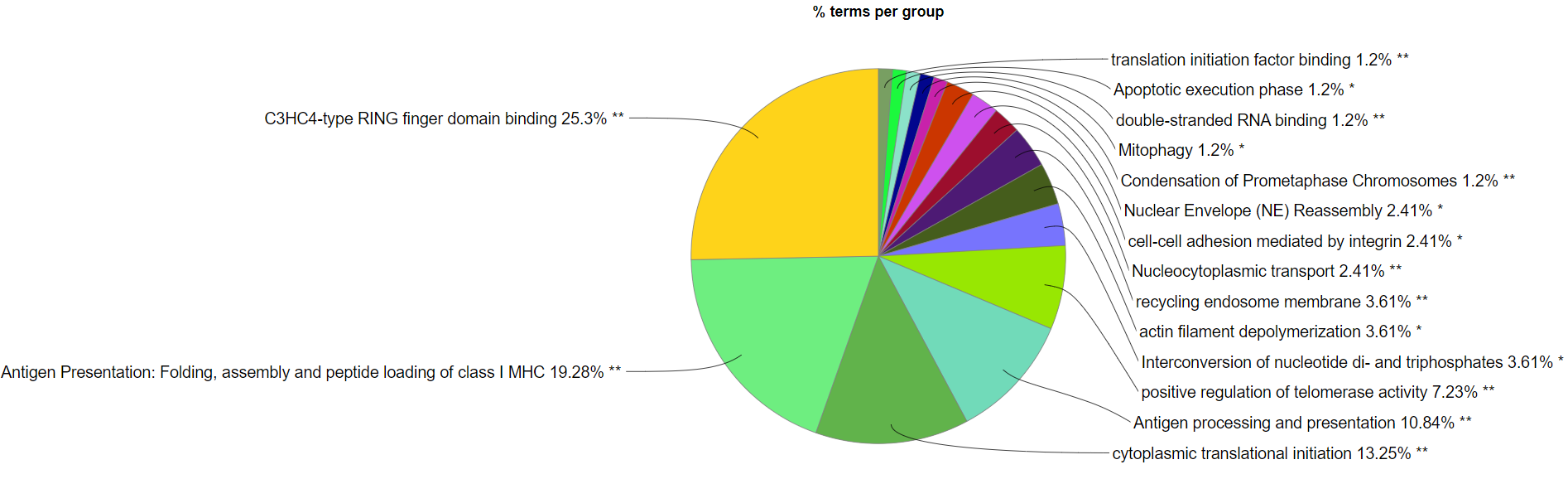

### 6-GSEA.pdf

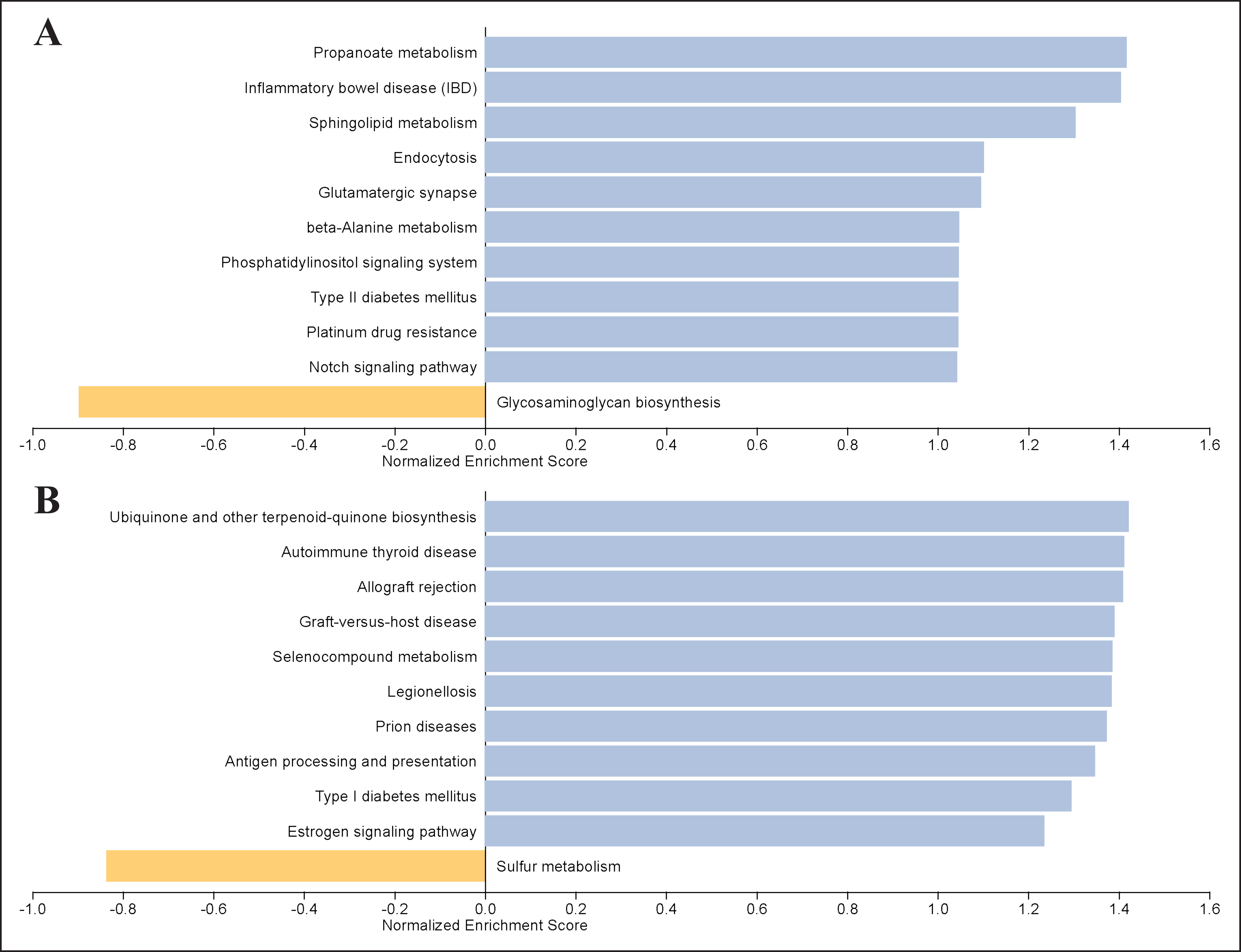

### 7_top_10-modified_bases_training_hela.png

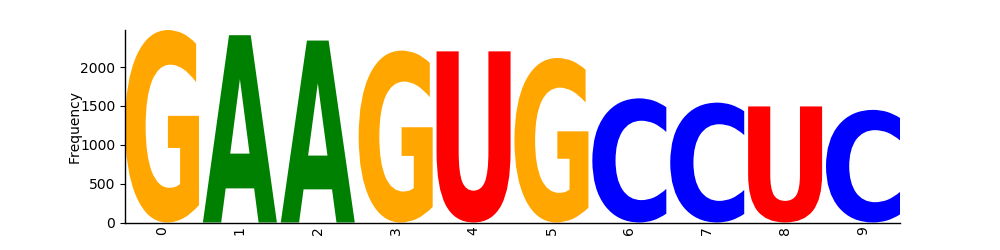

### 8_top_10-modified_bases_training_hek.png

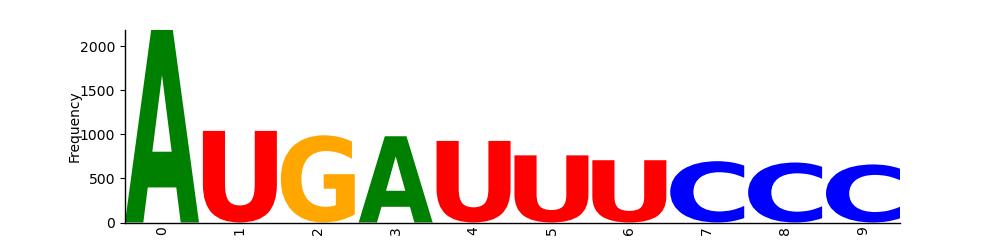
